# Soil chemistry determines whether defensive plant secondary metabolites promote or suppress herbivore growth

**DOI:** 10.1101/2021.05.14.444261

**Authors:** Lingfei Hu, Zhenwei Wu, Christelle A.M. Robert, Xiao Ouyang, Tobias Züst, Adrien Mestrot, Jianming Xu, Matthias Erb

## Abstract

Specialized metabolites mediate important interactions in both the rhizosphere and the phyllosphere. How this compartmentalized multifunctionality influences plant-environment interactions is unknown. Here, we investigated how the dual role of maize benzoxazinoids as leaf defenses and root siderophores shapes the interaction between maize and a major global insect pest, the fall armyworm. We find that benzoxazinoids suppress fall armyworm growth in soils with low bioavailable iron but enhance growth in soils with higher bioavailable iron. Manipulation experiments confirm that benzoxazinoids suppress herbivore growth under iron-deficient conditions but enhance herbivore growth when iron is present in its free form. This reversal of the protective effect of benzoxazinoids is not associated with major changes in plant primary metabolism. Plant defense activation is modulated by the interplay between soil iron and benzoxazinoids but does not explain fall armyworm performance. Instead, increased iron supply to the fall armyworm by benzoxazinoids in the presence of free iron enhances larval performance. This work identifies soil chemistry as a decisive factor for the impact of plant secondary metabolites on herbivore growth. It also demonstrates how the multifunctionality of plant secondary metabolites drives interactions between abiotic and biotic factors, with major consequences for plant health in variable environments.

## Introduction

Organismal traits are commonly co-opted for multiple functions (1–4). In complex and fluctuating environments, multifunctionality may lead to trade-offs, with important consequences for ecological and evolutionary dynamics (5–7).

Plant secondary (or specialized) metabolites are important mediators of species interactions in natural and agricultural systems (8). Many plant secondary metabolites have been documented to protect plants against insect herbivores by acting as toxins, digestibility reducers and / or repellents (9). Plant secondary metabolites also serve other functions; they can for instance act as signaling molecules (10), photoprotectants (11), antibiotics (12), soil nutrient mobilizers (13) and precursors of primary metabolites (14). Recent genetic work has highlighted that the same plant secondary metabolites may engage in multiple functions (4, 15, 16), leading to potentially important interactions between different environmental factors (6, 17). How this multifunctionality influences plant ecology and plant-insect interactions in complex environments is not well understood.

The soil environment can have a major impact on plant defense expression and plant-herbivore interactions. Soil nutrients and micronutrients can reprogram plant defenses, through cross-talk between defense and nutrient signaling (18, 19) or by influencing soil microbes which subsequently modulate plant defense responses (20, 21). In addition, soil nutrients can also influence plant-herbivore interactions by changing the nutritional value of the plant to herbivores (22). Whether plant secondary metabolites with dual functions in the rhizosphere and phyllosphere can mediate interactions between soil chemistry andherbivores remains unexplored, despite the potential ecological and agricultural importance of this phenomenon.

Benzoxazinoids are shikimic-acid derived secondary metabolites that are produced in high abundance by grasses such as wheat and maize. They evolved multiple times within the plant kingdom and are also found in various dicot families (23). Initially, benzoxazinoids were described as defense compounds that suppress and repel insect herbivores (24). Later genetic work revealed that benzoxazinoids also act as within-plant signaling compounds by initiating callose deposition upon pathogen and aphid attack (25, 26). Benzoxazinoids are also released into the rhizosphere in substantial quantities (27), where they can chelate iron (28), thus making it bioavailable (29). By consequence, benzoxazinoids can influence plant iron homeostasis. Recently, a link was documented between the iron chelating capacity and the interaction between maize plants and the western corn rootworm. This highly adapted insect is attracted by iron benzoxazinoid complexes and can use them for its own iron supply (29). Thus, it is conceivable that the multiple functions of benzoxazinoids may lead to trade-offs between their function as defenses and their functions as providers of essential micronutrients.

Here, we explore how the multifunctionality of benzoxazinoids shapes interactions between soil conditions and a leaf herbivore. By comparing soils that differ in their trace element composition, we uncover that the protective effect of maize benzoxazinoids against the fall armyworm can be reversed to a susceptibility effect in certain soils. Using micronutrient analyses and manipulative laboratory experiments, we document that this phenomenon can be fully explained by the interaction of benzoxazinoids with free iron in the soil. We further document that iron and benzoxazinoids interact to control leaf defenses, but that the benzoxazinoid-dependent susceptibility is best explained by increased iron supply to the fall armyworm. Taken together, these results provide a direct mechanistic link between soil properties and leaf-herbivore interactions and illustrate how plant secondary metabolite multifunctionality shapes plant-herbivore interactions.

## Results

### The effect of benzoxazinoids on herbivore performance depends on soil type

Benzoxazinoids increase leaf resistance to herbivores, but also interact with soil micronutrients and microbial communities in the soil (29–31). To test whether the defensive function of benzoxazinoids is modulated by soil properties, we collected soils from 8 different arable fields around Yixing (Jiangsu province, China), including both Anthrosols and Ferrosols (classification according to Chinese soil taxonomy, see Fig. S1 and Table S1 for basic soil characteristics). We then grew wild type B73 (WT) and benzoxazinoid-deficient *bx1* mutant plants in the different soils, infected them with fall armyworm larvae and measured plant performance, leaf damage and fall armyworm performance. On plants grown in Ferrosols, fall armyworm larvae gained more weight on *bx1* mutant plants than WT plants, as expected from the defensive function of benzoxazinoids (Fig. 1). However, on plants grown in Anthrosols, the larvae gained more weight on WT plants than *bx1* mutant plants (Fig. 1). Leaf damage did not differ between genotypes (Fig. S2), implying a change in the digestibility of the consumed leaf material. Overall, maize seedlings accumulated more biomass when growing in Anthrosols than Ferrosols, with no differences between genotypes (Fig. S2). To confirm that the herbivore growth patterns depend on benzoxazinoid biosynthesis, we tested additional *bx1* and *bx2* mutant alleles in the W22 background in an Anthrosol and a Ferrosol. Again, the larvae grew better on *bx1* and *bx2* mutant in the Ferrosol, but grew significantly less on the mutants in the Anthrosol (Fig. S3). Thus, the impact of benzoxazinoid biosynthesis on herbivore performance is strongly dependent on the soil type.

**Fig. 1.**
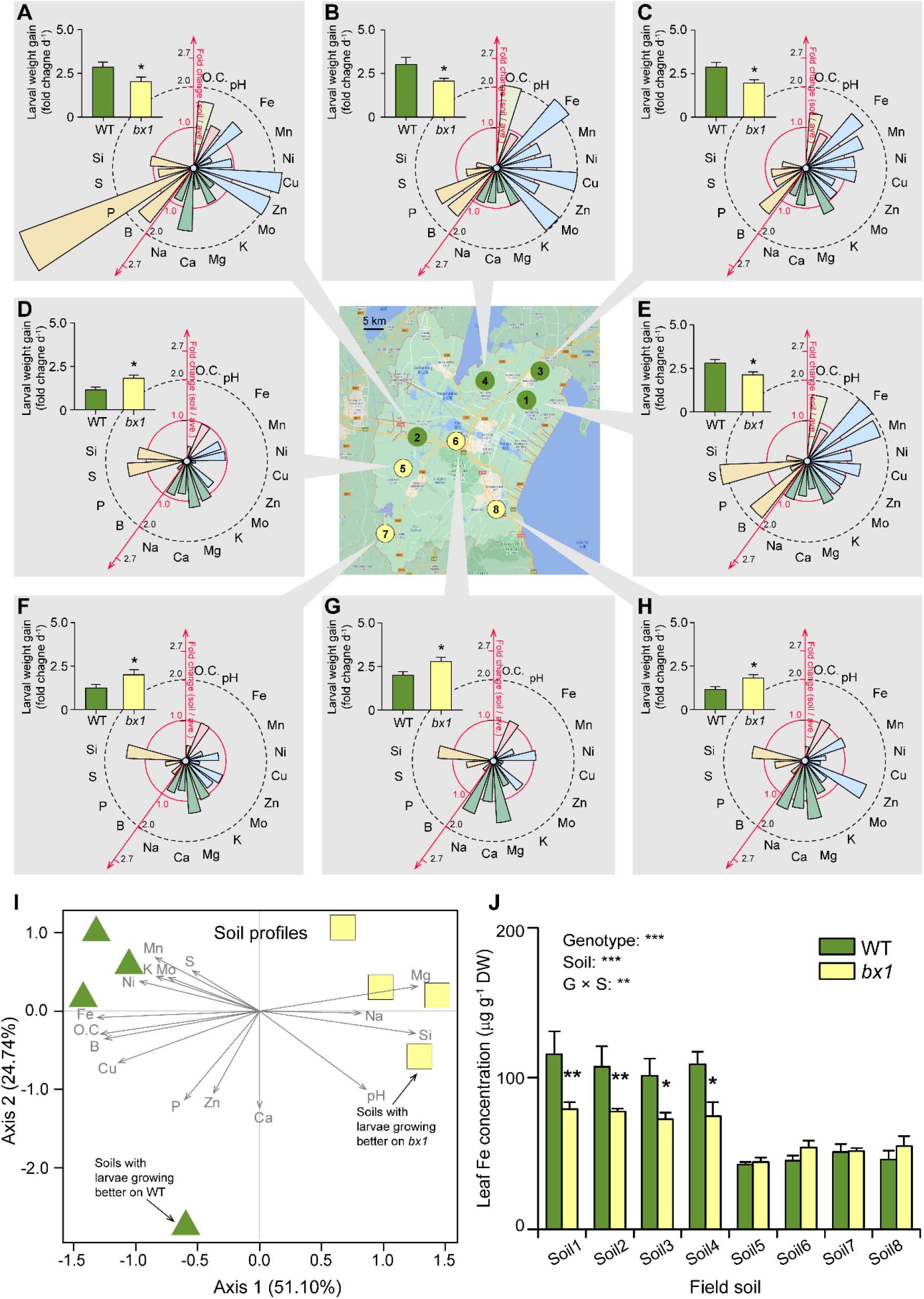
The effect of benzoxazinoids on herbivore performance depends on the soil type. Center: Map depicting soil collection sites around Yixing (China). Gray boxes (**A-H**): Growth of *Spodoptera frugiperda* caterpillars on wild type (WT) and benzoxazinoid deficient *bx1* mutant plants growing in the different soils (+SE, *n* = 10), together with respective soil properties. Soil properties are depicted as fold change relative to the average across all tested soils. See Fig. S1 for absolute values. Soils 1-4 are Anthrosols, soils 5-8 are Ferrosols. Asterisks indicate significant differences between plant genotypes (ANOVA; \**P* < 0.05). (**I**) Principal component analysis of field soil properties. Green triangles represent soils on which caterpillars grow better on WT plants. Yellow squares represent soils on which caterpillars grow better on *bx1* mutant plants. Vectors of soil parameters are shown as grey arrows. (**J**) Iron (Fe) contents in the leaves of WT and *bx1* plants grown in the different soils (+SE, *n* = 3, with 3 to 4 individual plants pooled per replicate). For full elemental analysis, refer to Fig. S4. DW, dry weight. O.C., organic carbon. Two-way ANOVA results testing for genotype and soil effects are shown (\*\**P* < 0.01; \*\*\**P* < 0.001). Asterisks indicate significant differences between genotypes within the same soil (pairwise comparisons through FDR-corrected LSMeans; \**P* < 0.05; \*\**P* < 0.01).

### Soil-dependent benzoxazinoid resistance is driven by root iron supply

Benzoxazinoids can chelate micronutrients and trace metals (28, 32), with the strongest quenching being observed for iron (29). We thus explored correlations between different soil properties and available micronutrients and the *bx1*-dependent impact on fall armyworm performance. Principal component analysis resulted in a clear separation of Anthrosols from Ferrosols (Fig. 1I). Ferrosols, on which larvae grew better on *bx1* than WT plants, had a slightly higher pH, less organic carbon, very low bioavailable iron, copper, boron and phosphorus, but higher sodium, magnesium and silicon than Anthrosols (Fig. 1I, Fig. S1). To further narrow down potential micronutrients that may drive the different genotype-specific performance of the fall armyworm, we screened micronutrient levels in the leaves of WT and *bx1* plants growing in the different soils. Analyses of variance revealed significant genotype effects for calcium and iron, with overall higher levels of both elements in the leaves of WT than *bx1* mutant plants (Fig. S4). Furthermore, a significant interaction between genotype and soil type was found for iron (Fig. 1J), with higher iron levels in WT plants than *bx1* mutant plants in Anthrosols with high available iron, and no difference in Ferrosols with very low available iron (Fig. 1J). Overall iron levels were enhanced in plants grown in Anthrosols compared to plants grown in Ferrosols (Fig. 1J), which is expected given that iron in Ferrosols is chiefly present as insoluble iron oxide. Quantitative analysis of the data revealed a strong association between higher iron levels in WT plants and higher fall armyworm performance (Fig. S5). A correlation was also observed for copper (Fig. S5), but this pattern was not associated with significant differences in copper levels between genotypes (Fig. S5). No correlation was observed for calcium (Fig. S5). The association between plant iron and fall armyworm performance was conserved in the W22 background (Fig. S6).

Based on these results, we hypothesized that differences in iron availability may determine the impact of benzoxazinoids on herbivore performance. To test this hypothesis, we grew WT and *bx1* mutant plants in nutrient solutions with different forms of iron (29). Plants where either grown in iron-deficient solutions (supplemented with NaCl or Na_2_SO_4_), solutions containing free, soluble iron (supplied as FeCl_3_ or Fe_2_(SO_4_)_3_) that requires chelation by siderophores for efficient uptake, or solutions containing a bioavailable iron complex (Fe-EDTA). Fall armyworm larvae grew generally better on benzoxazinoid-deficient *bx1* mutant plants, but effects depended on iron bioavailability and were reversed for plants in solutions containing free iron (Fig. 2A, Fig. S7). Leaf damage was similar across genotypes and iron treatments (Fig. 2B). Complementation of *bx1* mutant plants with pure DIMBOA added to the rhizosphere reverted the *bx1* phenotype in solutions containing free iron (Fig. 2C). These results show that the interaction between benzoxazinoids and iron availability can modulate the impact of benzoxazinoids on leaf herbivore performance.

**Fig. 2.**
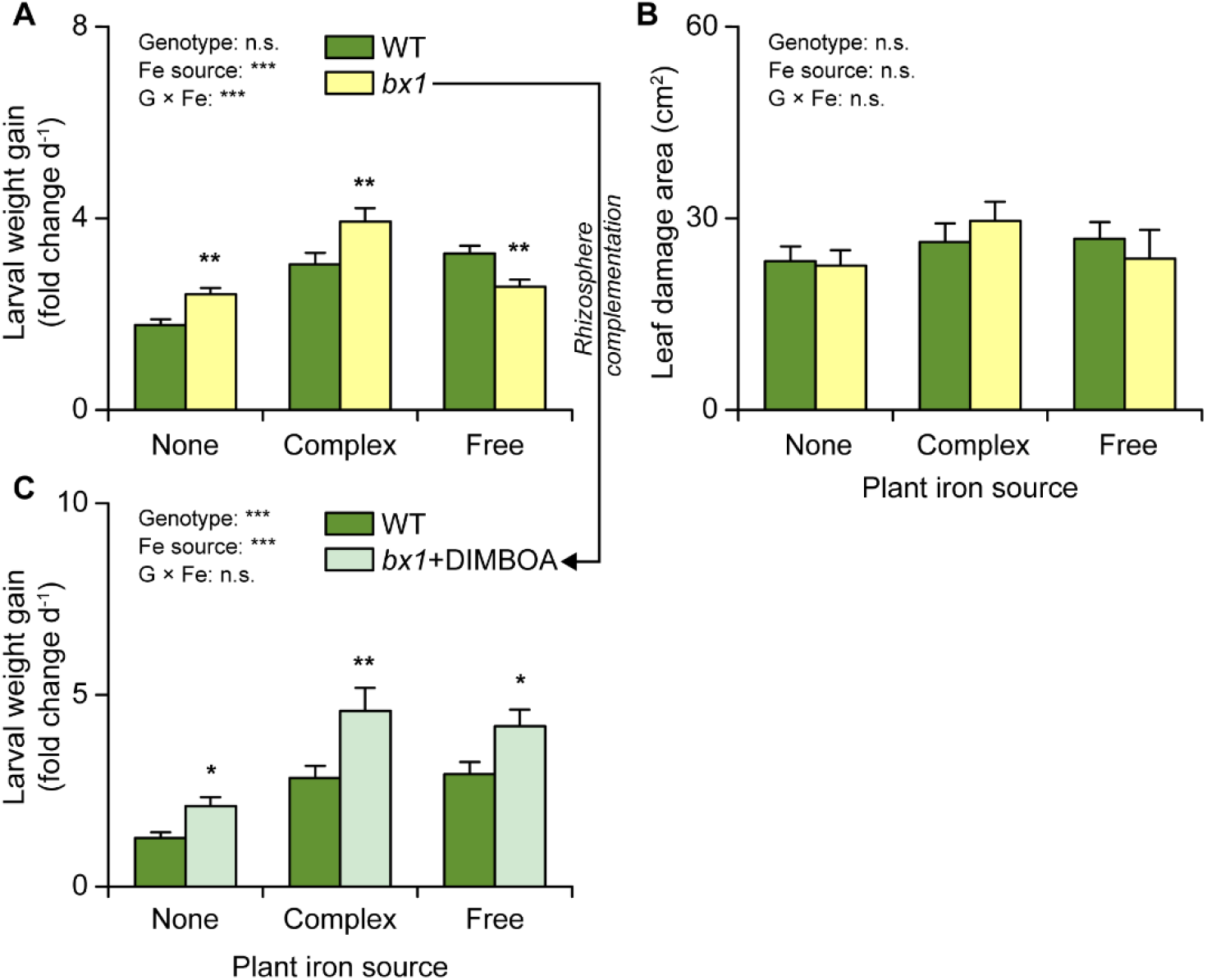
Soil-dependent benzoxazinoid resistance is driven by root iron (Fe) supply. (**A**) Growth of *Spodoptera frugiperda* feeding on WT and *bx1* plants supplied with different iron sources (+SE, *n* = 14-15). (B) Consumed leaf area (+SE, *n* = 14-15). (**C**) Growth of *S. frugiperda* feeding on WT and *bx1* plants complemented with pure DIMBOA added to the rhizosphere under different iron sources (+SE, *n* = 14-15). “None” nutrient solutions received either NaCl or Na_2_SO_4_. “Complex” nutrient solutions received Fe-EDTA. “Free” nutrient solutions received FeCl_3_ or Fe_2_(SO_4_)_3_. For full results showing genotype effects of all individual nutrient solutions, refer to Fig. S7. Two-way ANOVA results testing for genotype and iron source effects are shown (n.s. not significant; \*\*\**P* < 0.001). Asterisks indicate significant differences between genotypes within the same soil (pairwise comparisons through FDR-corrected LSMeans; \**P* < 0.05; \*\**P* < 0.01).

### Interactions between root iron supply and benzoxazinoids determine leaf iron homeostasis

*Bx1* mutant plants at the seedling stage are less efficient at taking up free iron than WT plants, and this trait is due to the absence of DIMBOA in the rhizosphere of *bx1* mutants (29). Thus, iron supply and benzoxazinoids likely interact to determine leaf iron homeostasis. In support of this hypothesis, we find that WT plants contain more iron in their leaves than *bx1* plants when grown in soils where iron is available in free or weakly complexed form, but not in soil where iron availability is low (Figs. 1, S4 and S7). To further explore this aspect, we measured the expression of genes involved in iron homeostasis in the leaves of WT and *bx1* mutant plants grown under different forms of iron supply (Fig. 3, Fig. S8). The tested genes included genes associated with iron transport such as *ZmYS1*, *ZmNRAMP1* and *ZmIRO2*, and genes that are likely involved in the biosynthesis and efflux of the mugineic acid family of siderophores such as *ZmRP1*, *ZmIDI4*, *ZmNAS3*, *ZmDMAS1* and *ZmTOM2* (33–35). We found strong interactions between iron availability and the *bx1* mutation for 7 of the 8 tested genes. Most iron homeostasis genes were highly expressed under iron-deficient conditions, and suppressed in the presence of complex iron, with no differences between WT and *bx1* mutant plants. When iron was present in its free form, most of these genes were strongly induced in the *bx1* mutant, but not in WT plants (Fig. 3). Exceptions to this pattern included *ZmNAS3*, which showed opposite expression patterns, and *ZmTOM2*, whose expression was not modulated by the *bx1* mutation (Fig. 3). These results show that root iron supply strongly modulates leaf iron homeostasis, with *bx1* mutants exhibiting iron deficiency gene expression patterns when grown in the presence of free iron.

**Fig. 3.**
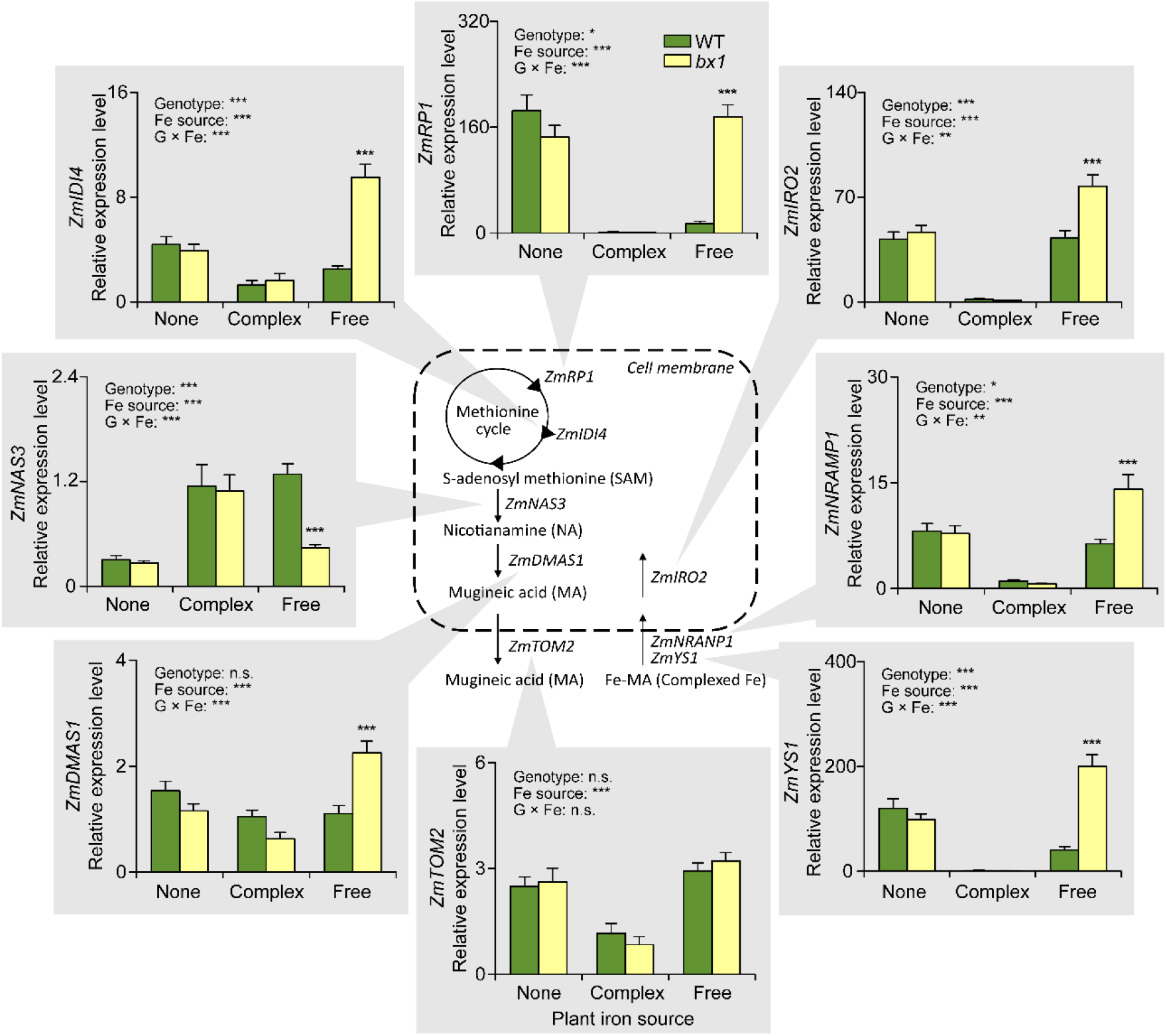
Interactions between root iron (Fe) supply and benzoxazinoids determine leaf iron homeostasis. Relative expression of genes involved in iron homeostasis in the leaves of WT and *bx1* mutant plants supplied with different iron sources (+SE, *n* = 7-8). “None” nutrient solutions received either NaCl or Na_2_SO_4_. “Complex” nutrient solutions received Fe-EDTA. “Free” nutrient solutions received FeCl_3_ or Fe_2_(SO_4_)_3_. For full results showing genotype effects of all individual nutrient solutions, refer to Fig. S8. Two-way ANOVA results testing for genotype and iron source effects are shown (n.s. not significant; \**P* < 0.05; \*\**P* < 0.01; \*\*\**P* < 0.001). Asterisks indicate significant differences between genotypes within the same soil (pairwise comparisons through FDR-corrected LSMeans; \*\*\**P* < 0.001).

### Changes in leaf herbivore performance are not explained by changes in leaf primary metabolism and defense expression

How can soil- and benzoxazinoid-dependent leaf iron homeostasis explain fall armyworm performance? In theory, iron homeostasis may indirectly affect leaf quality by changing leaf primary metabolism and defense expression (36–38), or directly by acting as a herbivore micronutrient (22). To test the first hypothesis, we measured soluble protein, hydrolysable amino acid and carbohydrate levels in the leaves of WT and *bx1* mutant plants grown under different iron regimes (Fig. 4, Fig. S9). No significant differences were found, suggesting that the different performance of the fall armyworm is not explained by major changes in leaf primary metabolites.

**Fig. 4.**
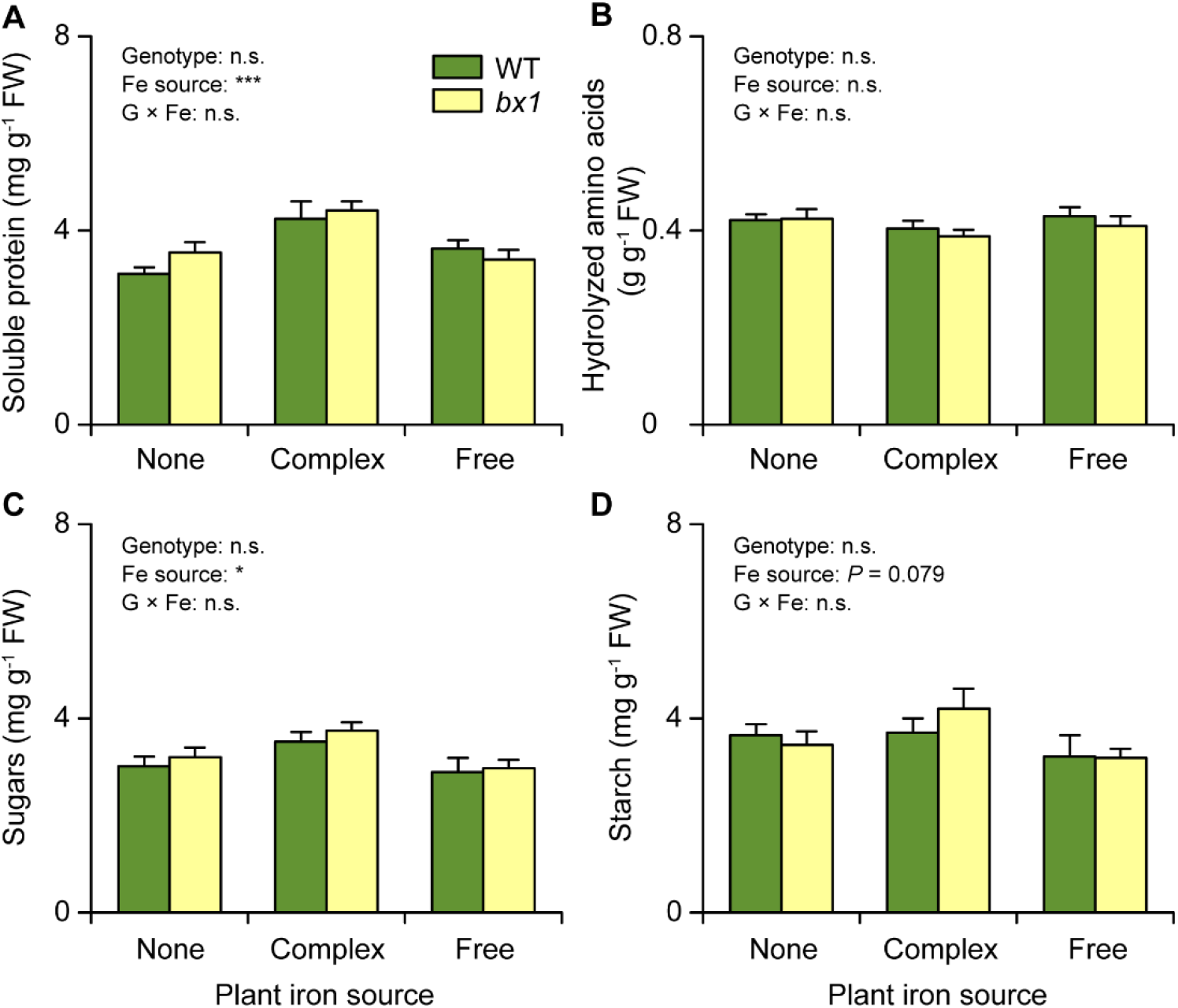
**Changes in leaf herbivore performance are not explained by changes in leaf primary metabolism**. Content of soluble protein (**A**), hydrolysable amino acids (**B**), sugars (**C**) and starch (**D**) in the leaves of wild type (WT) and *bx1* mutant plants supplied with different iron (Fe) sources (+SE, *n* = 14-15). “None” nutrient solutions received either NaCl or Na_2_SO_4_. “Complex” nutrient solutions received Fe-EDTA. “Free” nutrient solutions received FeCl_3_ or Fe_2_(SO_4_)_3_. For full results showing genotype effects of all individual nutrient solutions, refer to Fig. S9. Two-way ANOVA results testing for genotype and iron source effects are shown (n.s. not significant; \**P* < 0.05; \*\*\**P* < 0.001). No significant differences between genotypes within the same soil were observed (pairwise comparisons through FDR-corrected LSMeans).

Next, we measured the production of leaf defense metabolites that are produced independently of the benzoxazinoid biosynthesis pathway, including phenolic acids and the flavonoid maysin, a potential resistance factor against the fall armyworm (39), and the expression of defense marker genes, including a proteinase inhibitor (*ZmMPI*) (40) and a gene encoding the defense protein RIP2 (*ZmRIP2*), which is toxic to the fall armyworm *in vitro* (41). We detected significant interactions between iron availability and the *bx1* mutation for rutin and *ZmRIP2* expression (Fig. 5, Fig. S10). More rutin was produced in *bx1* mutant plants than WT plants grown in the presence of complex iron, but not when grown in iron-deficient and free iron solutions. *ZmRIP2* expression was lower in the *bx1* mutant than in WT plants, and these effects were most pronounced when plants were grown in iron-deficient and free iron solutions. These results show that root iron supply and benzoxazinoid biosynthesis interact to determine leaf-defense expression. At the same time, these interactions are unlikely to explain the observed differences in fall armyworm performance, as patterns do not match. *ZmRIP2* expression for instance was strongly reduced in the *bx1* mutant growing without iron or with iron in its free form, while fall armyworm performance was enhanced on *bx1* mutant plants grown without iron but suppressed in *bx1* plants grown with free iron.

**Fig. 5.**
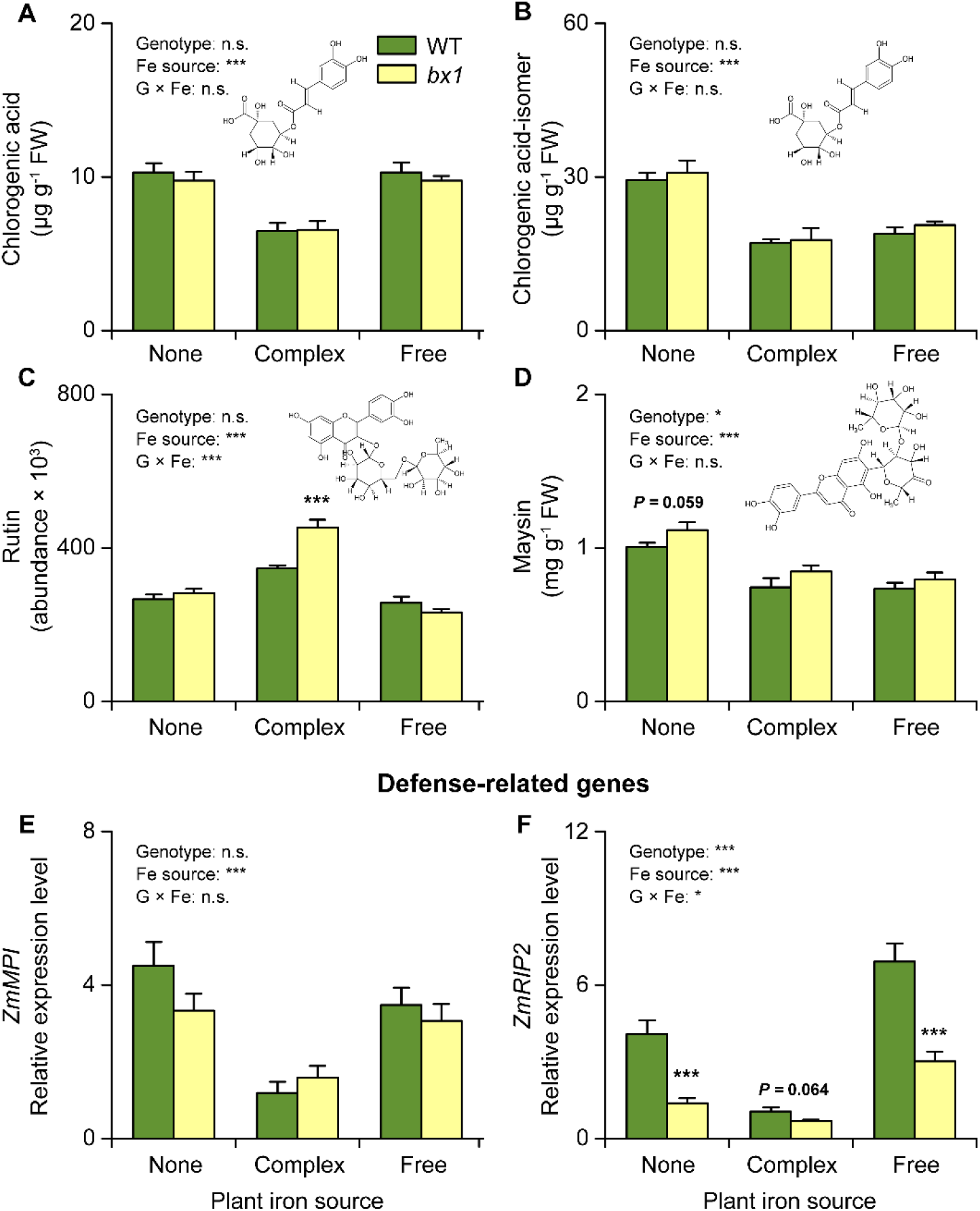
Soil iron (Fe) and benzoxazinoids interactively reprogram a subset of leaf defenses. (**A-D**) Concentrations of chlorogenic acid (A), chlorogenic acid-isomer (B), rutin (C) and maysin (D) in the leaves of WT and *bx1* mutant plants supplied with different iron sources (+SE, *n* = 8). FW, fresh weight. (**E** and **F**) Expression levels of *ZmMPI* (E) and *ZmRIP2* (F) in the leaves of WT and *bx1* plants supplied with different iron sources (+SE, *n* = 8). “None” nutrient solutions received either NaCl or Na_2_SO_4_. “Complex” nutrient solutions received Fe-EDTA. “Free” nutrient solutions received FeCl_3_ or Fe_2_(SO_4_)_3_. For full results showing genotype effects of all individual nutrient solutions, refer to Fig. S10. Two-way ANOVA results testing for genotype and iron source effects are shown (\**P* < 0.05; \*\*\**P* < 0.001). Asterisks indicate significant differences between genotypes within the same soil (pairwise comparisons through FDR-corrected LSMeans; \*\*\**P* < 0.001).

### Herbivore iron supply is associated with soil-dependent benzoxazinoid resistance

To test the hypothesis that benzoxazinoids may improve fall armyworm performance by supplying dietary iron, we first screened micronutrient concentrations in fall armyworm larvae fed on WT and *bx1* mutant plants grown in the different field soils. We found higher levels of iron in larvae feeding on WT than *bx1* mutant plants in soil with high iron availability. In soils with low iron availability, larval iron levels where low, and not different between plant genotypes (Fig. 6A). No significant effects were found for the other tested elements (Fig. S11). The same pattern for iron was observed for *bx1* and *bx2* mutants in the W22 genetic background (Fig. S12). Larvae feeding on plants growing in different iron solutions showed corresponding patterns, with significantly lower larval iron levels when feeding on *bx1* than WT plants grown together with free iron (Fig. 6B, Fig. S13). Adding DIMBOA to the rhizosphere of *bx1* mutant plants restored larval iron levels (Fig. 6C), thus providing a direct link between benzoxazinoids in the rhizosphere and larval iron homeostasis. To test whether fall armyworm performance is affected by iron supply, we measured larval growth in the iron transport deficient *ys1* mutant (33, 42). Larvae gained less weight in the *ys1* mutant compared to wild type B73 plants (Fig. 6D). Next, we conducted iron supplementation experiments by producing an iron-deficient diet and supplementing it with different forms of iron, including the DIMBOA iron complex Fe(III)(DIMBOA)_3_ at physiological concentrations. Fall armyworm larvae gained more weight when fed on iron supplemented diets, irrespective of the iron source (Fig 6E). At the tested concentration, DIMBOA alone had no negative effect on fall armyworm growth, which is in accordance with earlier results (30). From these experiments, we conclude that the interaction between soil micronutrient composition and benzoxazinoid biosynthesis directly influence iron homeostasis of a leaf herbivore, and that these effects can explain the soil-dependent impact of benzoxazinoids on herbivore performance.

**Fig. 6.**
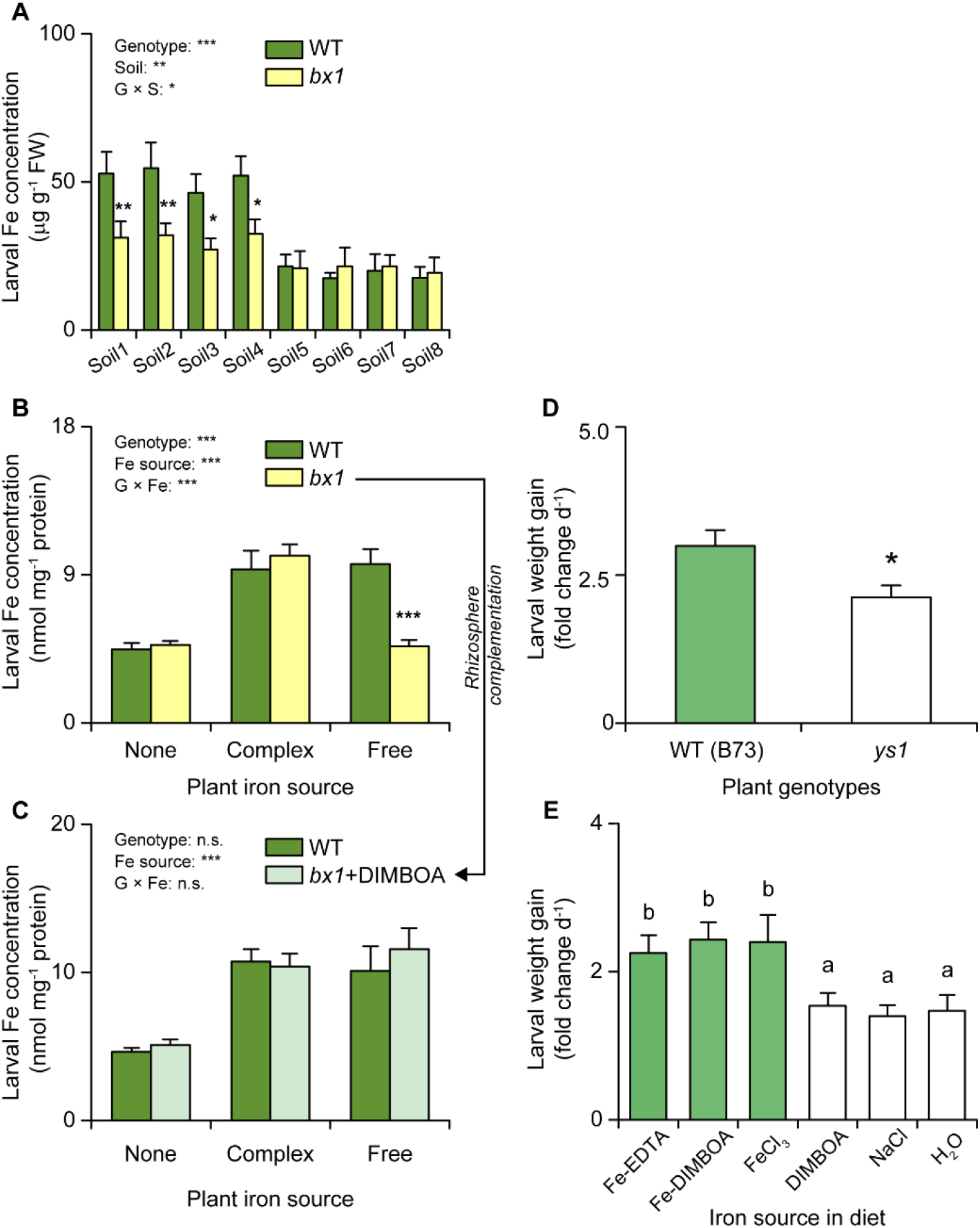
Herbivore iron (Fe) supply is associated with soil-dependent benzoxazinoid resistance. (**A**) Iron contents of *S. frugiperda* larvae feeding on WT and *bx1* plants grown in field soils (+SE, *n* = 3, with 3 to 4 individual larvae pooled per replicate). (**B** and **C**) Iron contents of *S. frugiperda* larvae feeding on WT and *bx1* plants (B), with DIMBOA rhizosphere complementation (C) under different iron source treatments (+SE, *n* = 5, with 3 individual larvae pooled per replicate). “None” nutrient solutions received either NaCl or Na_2_SO_4_. “Complex” nutrient solutions received Fe-EDTA. “Free” nutrient solutions received FeCl_3_ or Fe_2_(SO_4_)_3_. For full results showing genotype effects of all individual nutrient solutions, refer to Fig. S13. Two-way ANOVA results testing for genotype and iron source effects are shown (\*\*\**P* < 0.001). Asterisks indicate significant differences between genotypes within the same soil (pairwise comparisons through FDR-corrected LSMeans; \*\*\**P* < 0.001). (**D** and **E**) Growth of *S. frugiperda* larvae feeding on WT and *ys1* mutants (D, +SE, *n* = 13-16), and on iron-supplemented artificial diets (E, +SE, *n* = 12). Asterisks and letters indicate significant differences between genotypes or iron sources (one-way ANOVA followed by pairwise comparisons through FDR-corrected LSMeans; *P* < 0.05).

## Discussion

Organismal traits are commonly co-opted for multiple functions, which can lead to important context dependent performance patterns (5, 43). Here, we demonstrate that the multifunctionality of plant secondary metabolites results in conditional outcomes of plant-herbivore interactions, with soil properties determining whether the secondary metabolites promote or suppress leaf herbivore growth. Below, we discuss the mechanisms and agroecological implications of this finding.

Multifunctionality is a common property of plant secondary metabolites (16, 44, 45), with potentially important, but largely unexplored consequences for organismal interactions. We find that the protective effect of benzoxazinoids against an herbivore is determined and fully reversible by specific soil properties. A series of manipulative experiments in combination with the current state of knowledge (44, 46) allow us to infer the following scenario. When ingested by herbivores such as the fall armyworm, benzoxazinoids are rapidly deglycosylated (30). While the more stable forms such as DIMBOA can be detoxified through stereoselective reglycosylation (47), less stable aglucones such as HDMBOA can form reactive hemiacetals that form covalent bonds with thiol groups and can thus act as digestibility reducers as well as behavioral modulators (30, 48). All of these effects likely contribute to reduced herbivore weight gain of the fall armyworm in the presence of benzoxazinoids. At the same time however, benzoxazinoids are also released into the rhizosphere, where they interact with soil microbes (49) and effectively chelate free (29) and weakly bound iron, thus making it available to the plant. The higher iron uptake increases leaf iron levels, which benefits herbivores whose growth is limited by iron supply. The net impact of benzoxazinoid biosynthesis on the interaction between maize and leaf herbivores is thus governed by the strength of the negative effects of benzoxazinoids as digestibility reducers and the strength of the positive effects of benzoxazinoids as siderophores. By consequence, soil chemistry can tip the balance and determine whether benzoxazinoids have a net positive or negative effect on herbivore performance.

Plant nutrients in general (19, 50–54) and soil iron supply in particular (20, 21, 55), are increasingly recognized as important modulators of plant defense expression. In *Arabidopsis thaliana* for instance, the coumarin scopoletin is secreted under iron-deficiency and influences root microbiome assembly (21), likely including microbes that subsequently trigger systemic resistance in the plant by activating or priming hormonal defense pathways (20). In our work, we find that benzoxazinoid biosynthesis interacts with root iron supply to regulate a subset of leaf defenses, including the phenolic acid rutin and mRNA levels of the defense protein ZmRIP2. These effects are unlikely to be caused by changes in primary metabolism via leaf iron supply, as leaf carbohydrates and amino acids do not show any differences between treatments at this time. Interestingly, patterns of defense expression and leaf iron homeostasis markers also do not correspond: While iron homeostasis is most strongly affected by benzoxazinoids in the presence of free iron, defense expression is most strongly affected by the presence of the iron complex Fe-EDTA in the growth medium. As benzoxazinoids shape the rhizosphere microbiome (49, 56, 57), and as this effect is likely modulated by competition for iron, it is well possible that the type of iron source that is present in the growth solution has an impact on benzoxazinoid-microbiome interactions, which again will affect the activation of leaf defenses through systemic signaling. Further experiments will be required to explore this hypothesis. Although the observed modulation of leaf defenses may have contributed to fall armyworm growth, they are not directly responsible for the observed benzoxazinoid-dependent patterns, as benzoxazinoids affect fall armyworm growth differently in no iron and free iron treatments, without any change in benzoxazinoid-mediated defense expression. Thus, we infer that the direct effects of iron uptake on herbivore performance override potential indirect effects via root microbial communities or iron-dependent defense regulation.

Plant-herbivore interactions play an important role in shaping ecological communities and agricultural productivity (58, 59). Understanding the role of plant secondary metabolites as resistance factors is thus important for both fields. Our work shows that the suppressive effect of benzoxazinoids on fall armyworm growth, which likely contributes to plant resistance, fully depends, and can even be reversed, depending on soil characteristics. From an ecological point of view, soil and plant chemistry likely interact to determine the outcome of plant-herbivore interactions above ground. In this way, soil properties may determine plant and herbivore community composition even more strongly and dynamically than hitherto anticipated (60). From an agricultural point of view, the uncovered dependencies limit the use of benzoxazinoids as natural defenses against the fall armyworm, a global invasive pest that is currently threatening maize production in Africa and China. The finding that benzoxazinoids do not suppress the growth of the fall armyworm when growing certain soils is of particular importance in the context of the rapidly expanding maize production in Asia, as it shows the limits of using a core innate defense mechanism to broadly protect agroecosystems from an important invasive pest.

## Materials and Methods

### Plants and insects

The maize (*Zea mays* L.) genotypes B73 (referred to as WT), *bx1*/B73 (referred to as *bx1*) (31), W22, *bx1*/W22 (*bx1*::Ds), *bx2*/W22 (*bx2*::Ds) (61) and *ys1* (34, 42) were used in this study. Fall armyworm *Spodoptera frugiperda* (J.E. Smith, 1791) larvae were reared on artificial diet as described previously (62).

### Plant and herbivore performance in field soils

To determine the impact of available soil nutrients on plant performance and herbivore resistance, we collected 8 soils from different arable fields in Yixing, Jiangsu province, China (Table S1). The soils were firstly air-dried and then individually passed through 2 mm sieve, homogenized, and used to fill 200 mL pots (11 cm depth and 5 cm diameter). B73, *bx1*/B73, W22, *bx1*/W22 and *bx2*/W22 plants were then individually grown in these soils. Pots were randomly placed on a greenhouse table (26 °C ± 2 °C, 55% relative humidity, 14:10 h light/dark, 50,000 lm m^-2^) and rearranged weekly. Plants were watered once a week. 16 days after planting, we measured the shoot dry weight, leaf chlorophyll content, elemental composition (*n* = 3, with 3-4 plants pooled per replicate) and larval growth (*n* = 10) on each maize plant (see below for details). Chlorophyll contents were determined using a SPAD-502 meter (Minolta Camera Co., Japan).

### Plant and herbivore performance in nutrient solutions

To assess the impact of different forms of iron on plant performance, a soil-free growth system was used as described previously (29). Briefly, B73 and *bx1* seeds were individually wrapped in two layers of paper. The paper rolls with the seeds were put in 200 mL pots (11 cm depth and 5 cm diameter). Pots were supplied with 40 mL Milli-Q water, covered with aluminum (Al) foil and then placed in the greenhouse (26°C ± 2°C, 55% relative humidity, 14:10 h light/dark, 50,000 lm m^-2^). One week after the start of germination, the remaining seed shell was removed from the germinating seedlings to reduce the influence of residual iron in the endosperm. Plants were then grown in nutrient solutions containing complexed iron (Fe-EDTA), free iron [FeCl_3_, Fe_2_(SO_4_)_3_] or no iron (NaCl, Na_2_SO_4_) sources. For a complete description of the nutrient solution, see (29). The final concentrations of the different forms of iron in the solution were 250 µM Fe-EDTA, 250 µM FeCl_3_ or 125 µM Fe_2_(SO_4_)_3_. The respective Fe-free control solutions contained 750 µM NaCl or 375 µM Na_2_SO_4_ to control for effects of Cl^-^, SO_4_^2-^ and Na^+^ in the iron salt treatments. The pH of the nutrient solutions was adjusted to 5.5 using KOH. All the chemicals were bought from Sigma (Sigma Aldrich, Beijing, China). Three weeks after germination, we quantified the gene expression, primary and secondary metabolites of the youngest fully developed leaf (*n* = 8), and larval growth on each plant (*n*= 14-15), as described below.

To determine whether DIMBOA is sufficient to restore the resistance of *bx1* plants to those of WT plants, WT and *bx1* plants were treated as described above. One week after germination, the plants were supplied with nutrient solutions containing Fe-EDTA, FeCl_3_, or NaCl. The nutrient solution for the *bx1* mutants was complemented with 300 µg DIMBOA, which corresponds to physiological concentration of DIMBOA that accumulate in the rhizosphere of B73 plants (29). The larval growth on each plant were then assessed as described below (*n* = 15).

To investigate the connections between plant Fe acquisition and larval growth, B73 and *ys1* plants were treated as described above (*n* = 13-16). One week after germination, the plants were grown in nutrient solutions supplied with EDTA-Fe. Three weeks after germination, larval growth on each plant was recorded.

### Soil, plant and herbivore nutrient analyses

To characterize nutrients in bulk field soil, soil samples were air-dried and then individually ground and passed through 1 mm sieve. The available Fe, Mn, Ni, Cu and Zn were extracted according to China Environmental Protection standards (HJ 804-2016). Briefly, 10.0 g soil sample was mixed with 20 mL extraction buffer (0.005 M diethylenetriaminepentaacetic acid [DPTA], 0.01 M CaCl_2_, 0.1 M triethanolamine [TAE], pH = 7.3), and then shaken (180 r/min) for 2 h at 20 °C. After shaking, the mixture was centrifuged, and supernatant was collected, and the concentrations of Fe, Mn, Ni, Cu and Zn were determined by inductively coupled plasma-mass spectrometry (ICP-MS) (NexION300X, PerkinElmer, USA). The parameters used during the ICP-MS measurements were: RF generator power output: 1600 W; argon flows: plasma, 1.5 L min^−1^; nebulizer: 1.09 L min^−1^, KED gas: helium, at flow 3.5 mL min^−1^; optimization on masses of ^9^Be, ^24^Mg, ^115^In, ^238^U, ^140^Ce; data acquisition: dwell time of 50 ms, 3 points per peak, acquisition time of 3 s. ^57^Fe, ^55^Mn, ^60^Ni, ^63^Cu, and ^66^Zn were used as analytical masses to reduce interferences. A 40 μg L^-1^ Rh solution as an internal standard in order to compensate any possible signal instability, and a washing cycle of at least 30 s was settled between two subsequent samples with the aim to eliminate any memory effects. All reported data were blank corrected. In order to monitor constantly the overall accuracy level of the method, a blank was run up every eight samples. A standard reference material, CRM Cabbage, GBW10014 (GSB-5), prepared by the Institute of Geophysical and Geochemical Exploration of China, was run every twelve samples to determine the accuracy of the analytical methods. The mixed standard samples (PerkinElmer, cat. no. N9300233) from 1 µg L^-1^ to 200 µg L^-1^ were used to build a standard curve, with correlation coefficient (R^2^) of each element being higher than 0.999. The absolute quantities of Fe, Mn, Ni, Cu and Zn were calculated according to the standard curve.

Soil pH, NH_4_^+^, available P, S, Si, B, Mo and exchangeable K^+^, Na^+^, Ca^2+^, Mg^2+^ were extracted and determined according to the China National Standard Methods. Briefly, soil pH was determined in a 2.5:1 water/soil suspension using a pH meter (LY/T1239-1999). NH_4_^+^ was measured using alkali hydrolysis diffusion (LY/T 1231-1999). Available P was determined by hydrochloric acid and ammonium fluoride (LY/T 1233–1999). Available S was extracted with calcium phosphate-acetic acid and quantified with quantified with turbidimetric method using barium sulfate (LY/T 1265-1999). Available Si was extracted by sodium acetate and quantified by the silicon-molybdenum blue colorimetry (LY/T 1266-1999). Available B was extracted by boiled deionized water and determined by the azomethine-H spectrophotometric method (LY/T 1258-1999). Available Mo was extracted by acid-ammonium-oxalate and quantified by colorimetry using potassium thiocyanate (LY/T 1259-1999). Exchangeable K^+^, Na^+^, Ca^2+^ and Mg^2+^ were extracted by ammonium acetate and determined by flame photometry (LY/T 1246-1999) and atomic absorption spectrophotometry (LY/T 1245-1999), respectively.

For plant micronutrient analyses, plant leaves were oven dried. Three or four individual plants were pooled as one replicate. The samples were digested in 6 ml of 15 M HNO_3_ and 10 M H_2_O_2_ at 190℃ for 35 min with MARS 6 CLASSIC (CEM Corp., Matthews, NC, USA) as described (63). After digestion, the samples were dissolved in deionized water. The concentrations of Mg, Fe, Mn, Ni, Cu and Zn were determined by ICP-MS as described above. The concentrations of K, Ca, Na, P, Si, B and Mo were quantified by the ICP-optical emission spectrometric method according to the China National Standard Method (GB/T 35871-2018).

For micronutrient analyses of fall armyworm larvae, three or four larvae were pooled together as one replicate. The elements were extracted and determined as described above.

### Herbivore growth and damage assays

To assess the *S. frugiperda* growth on maize plants, individual starved and pre-weighted second instar larva were introduced into cylindrical mesh cages (1 cm height and 2.5 cm diameter). The cages were then clipped onto the leaves of maize plants (one cage per plant). The position of each cage was moved regularly to provide sufficient food supply for the larvae. Larval weight was recorded 5 days after the start of the experiment. The remaining leaves were scanned, and the removed leaf area was quantified using Digimizer 4.6.1 (Digimizer).

### Primary metabolite analyses

Soluble protein was extracted and quantified using a Bradford assay (*n* = 8) (64). Amino acids were hydrolyzed and quantified by UHPLC-MS (Waters, USA) according to a previously published protocol (*n* = 8) (65). Starch and soluble sugars (glucose, sucrose, and fructose) were extracted and quantified as described previously (*n* = 8) (66).

### Secondary metabolite analyses

To quantify the influences of Fe forms on secondary metabolites, three maize plants were pooled, homogenized, and ground by liquid nitrogen (*n* = 4 pools per Fe treatment). Seventy mg of ground samples were extracted in 700 μL of acidified H_2_O/MeOH (50:50 v/v; 0.1% formic acid), and then analyzed with an Acquity UHPLC system coupled to a G2-XS QTOF mass spectrometer (MS) equipped with an electrospray source (Waters, USA) as described (49). Briefly, compounds were separated on an Acquity BEH C18 column (2.1×50 mm i.d., 1.7 μm particle size). Water (0.1% formic acid) and acetonitrile (0.1% formic acid) were employed as mobile phases A and B. The elution profile was: 0-3.50 min, 99-72.5% A in B; 3.50-5.50 min, 72.5-50% B; 5.51-6.50 min 100% B; 6.51-7.51 min, 99% A in B. The flow rate was 0.4 mL/min. The column temperature was maintained at 40°C, and the injection volume was 1 μL. The QTOF MS was operated in negative mode. The data were acquired over an *m*/*z* range of 50-1200 with scans of 0.15 s at collision energy of 4 V and 0.2 s with a collision energy ramp from 10 to 40 V. The capillary and cone voltages were set to 2 kV and 20 V, respectively. The source temperature was maintained at 140°C, the desolvation was 400°C at 1000 L h^-1^ and cone gas flows was 50 L/h. Accurate mass measurements (< 2 ppm) were obtained by infusing a solution of leucin encephalin at 200 ng/mL at a flow rate of 10 mL/min through the Lock Spray probe (Waters). The abundance of chlorogenic acid, chlorogenic acid-isomer, rutin were determined based on peak areas. The absolute quantities of maysin were determined using standard curves obtained from purified maysin as described (67).

### Gene expression analyses

Quantitative real time PCR (QRT-PCR) was used to quantify gene expressions. Total RNA was isolated from maize leaves using the GeneJET Plant RNA Purification Kit (Thermo Scientific, USA) following the manufacturer’s instructions (*n* = 8). Three hundred nanograms of each total RNA sample was reverse transcribed with SuperScript® II Reverse Transcriptase (Invitrogen, USA). The QRT-PCR assay was performed on the LightCycler® 96 Instrument (Roche, Switzerland) using the KAPA SYBR FAST qPCR Master Mix (Kapa Biosystems, USA). The maize actin gene *ZmActin* was used as an internal standard to normalize cDNA concentrations. The relative gene expression levels of target genes were calculated using the 2^-ΔΔCt^ method (68). The primers of all tested genes are provided in Table S2.

### Larval iron analyses

To determine iron concentrations in *S. frugiperda* larvae, three larvae were pooled and homogenized (*n* = 5 pools). Total protein of the larval lysates was extracted with 200 µL lysis buffer (20 mM Tris, 137 mM NaCl, 1% Triton X-100, 1% glycerol). The concentrations of extracted protein were quantified with a Bradford assay, and then denatured using a described protocol (69). After denaturation, 50 µL of the supernatant were taken for iron measurements using the Iron Assay Kit (Sigma, USA) following the manufacturer’s instructions, with the modification that 25 µL of saturated ammonium acetate were added and mixed to adjust the pH before measuring absorbance at 593 nm. The absolute quantities of iron were determined using standard curves made from pure FeCl_3_ according to the manufacturer’s instructions.

### Larval growth on diets with exogenous iron sources

To evaluate the direct effect of iron on *S. frugiperda* growth, we prepared the artificial diets containing different iron sources according to the methods as described by (70), with some modifications. Briefly, 17 g of agar were dissolved in 500 mL of water at 50°C and mixed with 5 g dried leaf material of *bx1* plants, 25 g casein, 23 g sucrose, 12 g yeast extract, 9 g Wesson salt mixture, 3.5 g ascorbic acid, 2.5 g cholesterol, 1.5 g sorbic acid, 5 mL raw linseed oil, 1.5 mL formalin and 9 mL vitamin mixture (100 mg nicotinic acid, 500 mg riboflavin, 233.5 mg thiamine, 233.5 mg pyridoxine, 233.5 mg folic acid and 20 mg L^-1^ biotin in water). Fe-EDTA, Fe(III)(DIMBOA)_3_, FeCl_3_, DIMBOA, NaCl, or H_2_O was then added to the diet at final concertation 50 μM, which corresponds to physiological concentration in maize xylem sap (29). The produced diet was aliquoted into solo cups. One starved and pre-weighted second instar larva was individually introduced into the solo cups. Diets were replaced every other day. Larval weight was recorded 5 days after the start of the experiment (*n* = 12).

### Statistical analyses

Larval growth, leaf damage, gene expression, and metabolite data were analyzed by analysis of variance (ANOVA) followed by pairwise or multiple comparisons of Least Squares Means (LSMeans), which were corrected using the False Discovery Rate (FDR) method (71). Normality was verified by inspecting residuals, and homogeneity of variance was tested through Shapiro-Wilk’s tests using the “plotresid” function of the R package “RVAideMemoire” (72). Datasets that did not fit assumptions were natural log-transformed to meet the requirements of equal variance and normality. For the redundancy analysis (RDA), raw data were first scaled with the “scale” function in R. PCAs were then performed with the “MVA” function of “RVAideMemoire” package and the “rda” function of “vegan” package (72, 73). All statistical analyses were conducted with R 3.4.4 (R Foundation for Statistical Computing, Vienna, Austria) using the packages “car”, “emmeans”, and “RVAideMemoire” (72–75).

### Accession numbers

The sequence data of maize genes can be found in the MaizeGDB database under the following accession numbers *ZmActin* (GRMZM2G126010), *ZmMPI* (GRMZM2G028393), *ZmRIP2* (GRMZM2G119705), *ZmRPI* (GRMZM2G035599), *ZmIDI4* (GRMZM2G067265), *ZmNAS3* (GRMZM2G478568), *ZmDMAS1* (GRMZM2G060952), *ZmTOM2* (GRMZM5G877788), *ZmYS1* (GRMZM2G156599), *ZmNRAMP1* (GRMZM2G178190) and *ZmIRO2* (GRMZM2G057413).

## Acknowledgements

We thank Prof. Nicolaus von Wirén from IPK Gatersleben for sharing the *ys1* mutants, Dr. Xianwen Zhang from Zhejiang University for propagating the seeds. **Funding:** This work was supported by the National Natural Science Foundation of China (42007029), the Young Elite Scientists Sponsorship Program by the CAST (2019QNRC002), the Interfaculty Research Collaboration “One Health” of the University of Bern, and the Fundamental Research Funds for the Central Universities (2020QNA6009). **Data and materials availability:** All data needed to evaluate the conclusions in the paper are present in the paper and/or the supplementary materials. Additional data related to this paper may be requested from the authors.

## Supplementary Materials

**Fig. S1.**
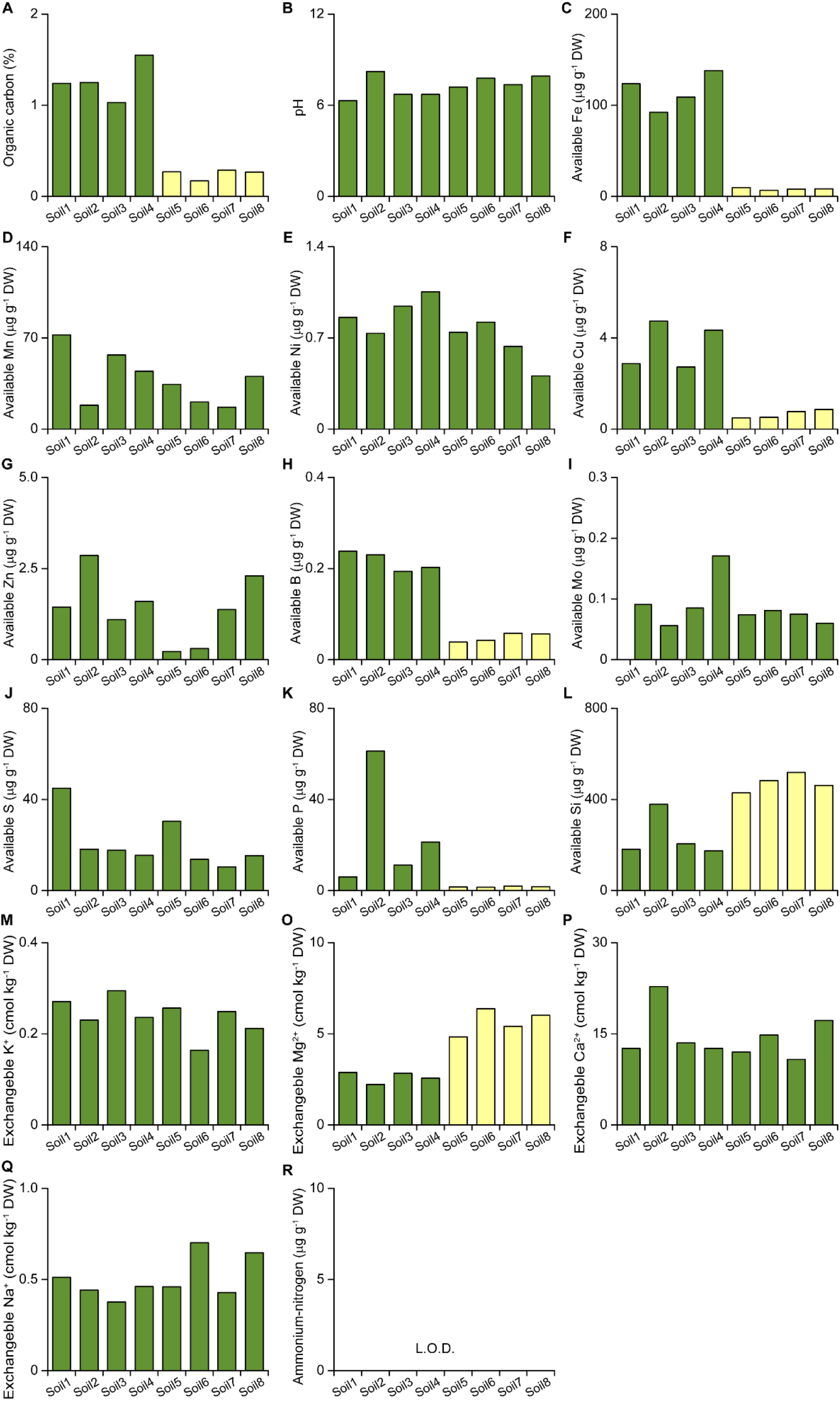
Soil nutrients and pH in different field soils. The content of organic carbon (A), pH (B), available iron (Fe, C), manganese (Mn, D), nickel (Ni, E), copper (Cu, F), zinc (Zn, G), boron (B, H), molybdenum (Mo, I), sulphur (S, J), phosphorous (P, K), silicon (Si, L), exchangeable potassium (K, M), magnesium (Mg, O), calcium (Ca, P), sodium (Na, Q) and ammonium-nitrogen (R) in field soils. DW, dry weight. L.O.D, below the limit of detection. Green bars correspond to soils where fall armyworm larvae grow better on wild type than *bx1* mutant plants. Yellow bars correspond to soils where fall armyworm larvae grow better on *bx1* mutant than wild type plants.

**Fig. S2.**
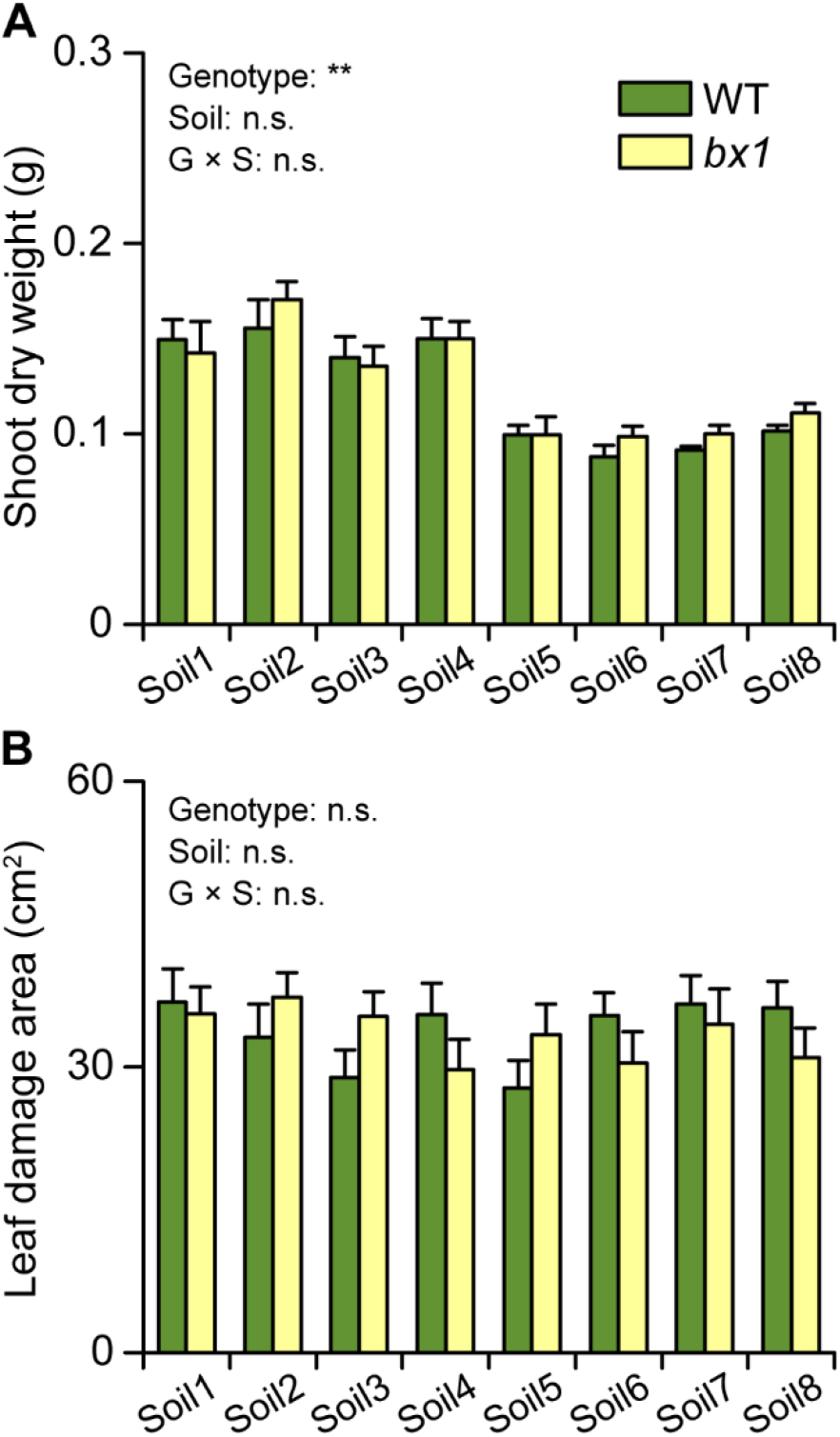
Plant performance and fall armyworm consumed leaf area. (A) Shoot dry weight (A) of wild type (WT) and *bx1* plants grown in field soils (+SE, *n* = 10). (B) Consumed leaf area by *Spodoptera frugiperda* larvae feeding on WT and *bx1* plants grown in field soils (+SE, *n* = 10). Two-way ANOVA revealed no significant soil or genotype effects (*P* > 0.05).

**Fig. S3.**
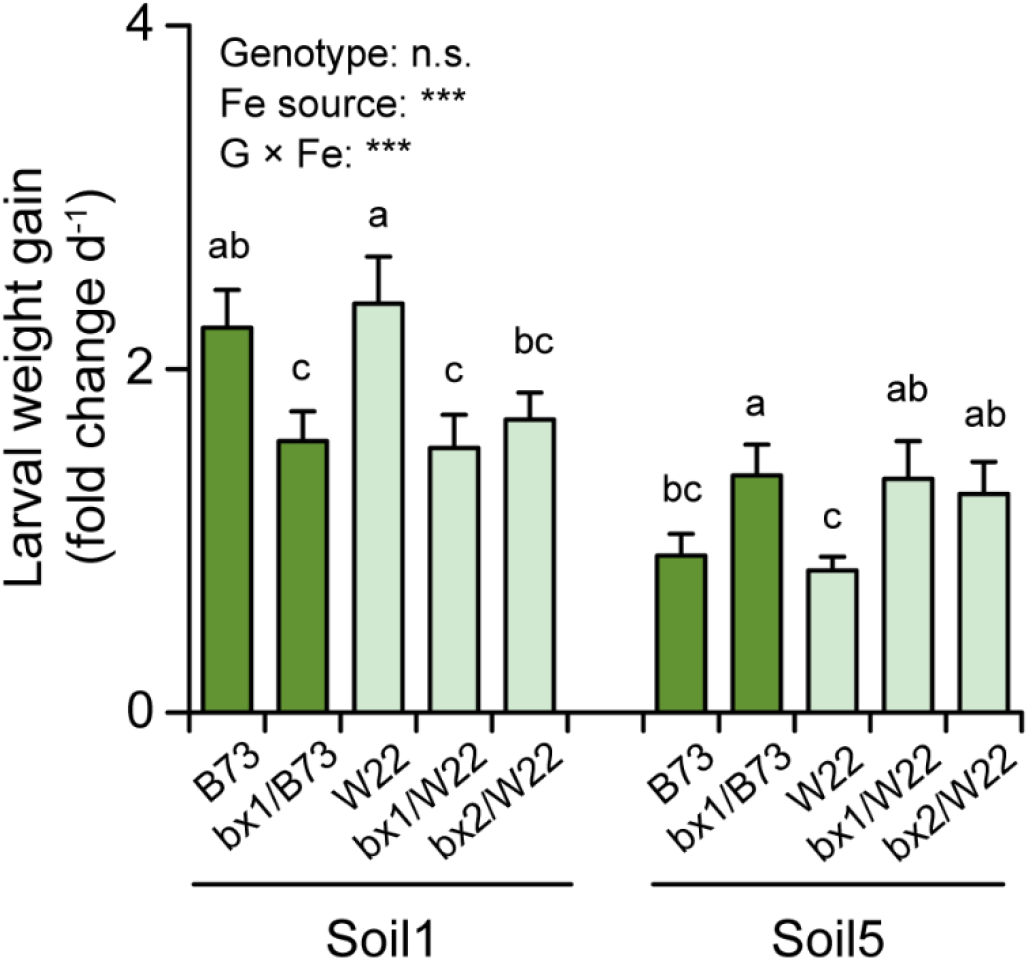
Soil type modulates the impact of benzoxazinoids on herbivore performance in the W22 genetic background. Growth of *S. frugiperda* larvae on maize plants carrying *bx1* and *bx2* mutant alleles in the W22 background in two of selected soils (+SE, *n* = 12). Results of two-way ANOVAs are shown (n.s. not significant; \*\*\**P* < 0.001). Different letters indicate significant differences between genotypes within the same soil (two-way ANOVA followed by pairwise comparisons through FDR-corrected LSMeans; *P* < 0.05).

**Fig. S4.**
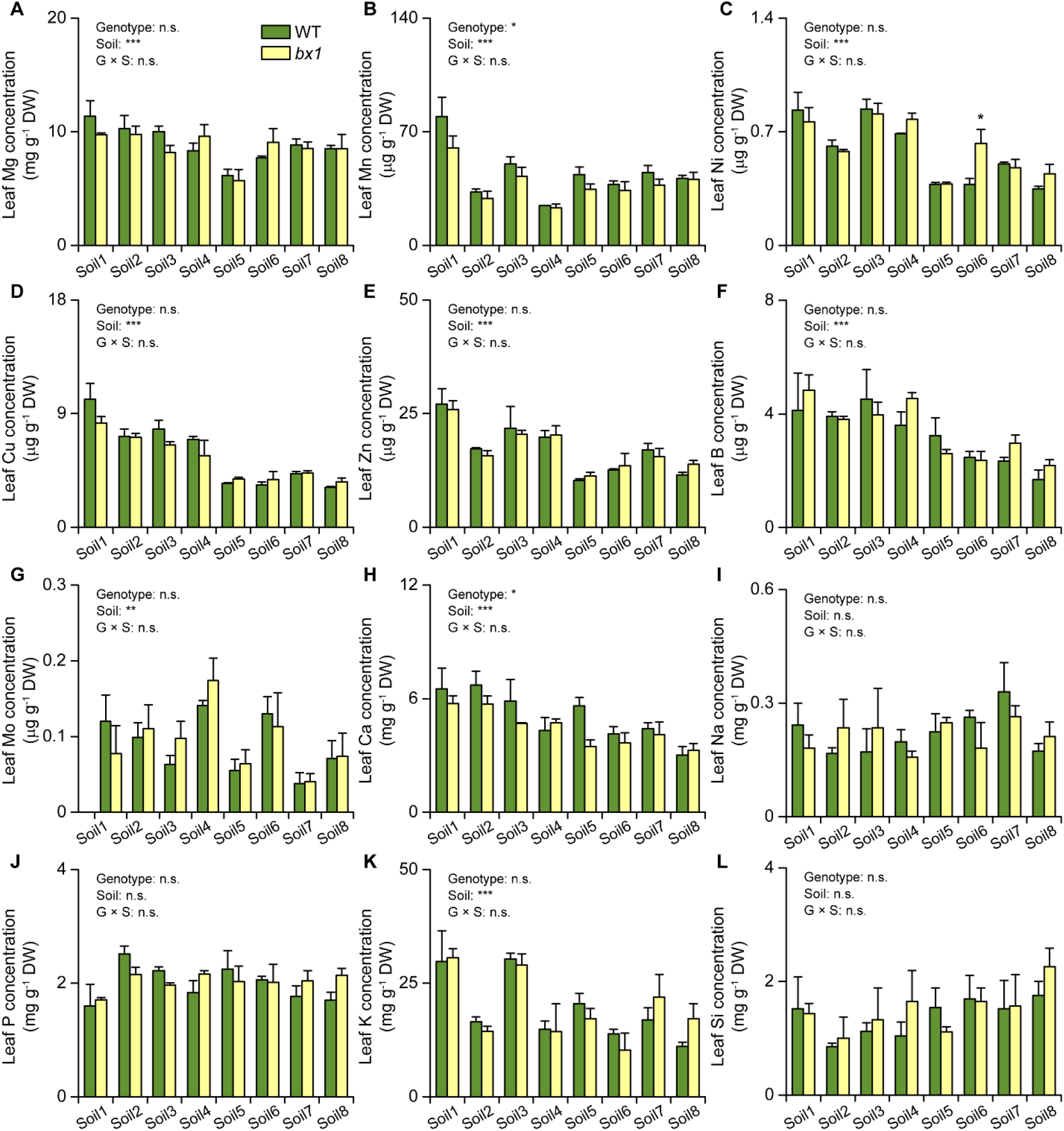
Leaf elemental analysis of wild type (WT) and *bx1* plants in different field soils. Concentrations of magnesium (Mg, A), manganese (Mn, B), nickel (Ni, C), copper (Cu, D), zinc (Zn, E), boron (B, F), molybdenum (Mo, G), calcium (Ca, H), sodium (Na, I), phosphorous (P, J), potassium (K, K), and silicon (Si, L) in WT and *bx1* plants grown in field soils (+SE, *n* = 3, with 3 to 4 individual plants pooled per replicate). DW, dry weight. Results of two-way ANOVAs are shown (n.s. not significant; \**P* < 0.05; \*\**P* < 0.01; \*\*\**P* < 0.001). Asterisks indicate significant differences between genotypes within the same soil (two-way ANOVA followed by pairwise comparisons through FDR-corrected LSMeans; \**P* < 0.05).

**Fig. S5.**
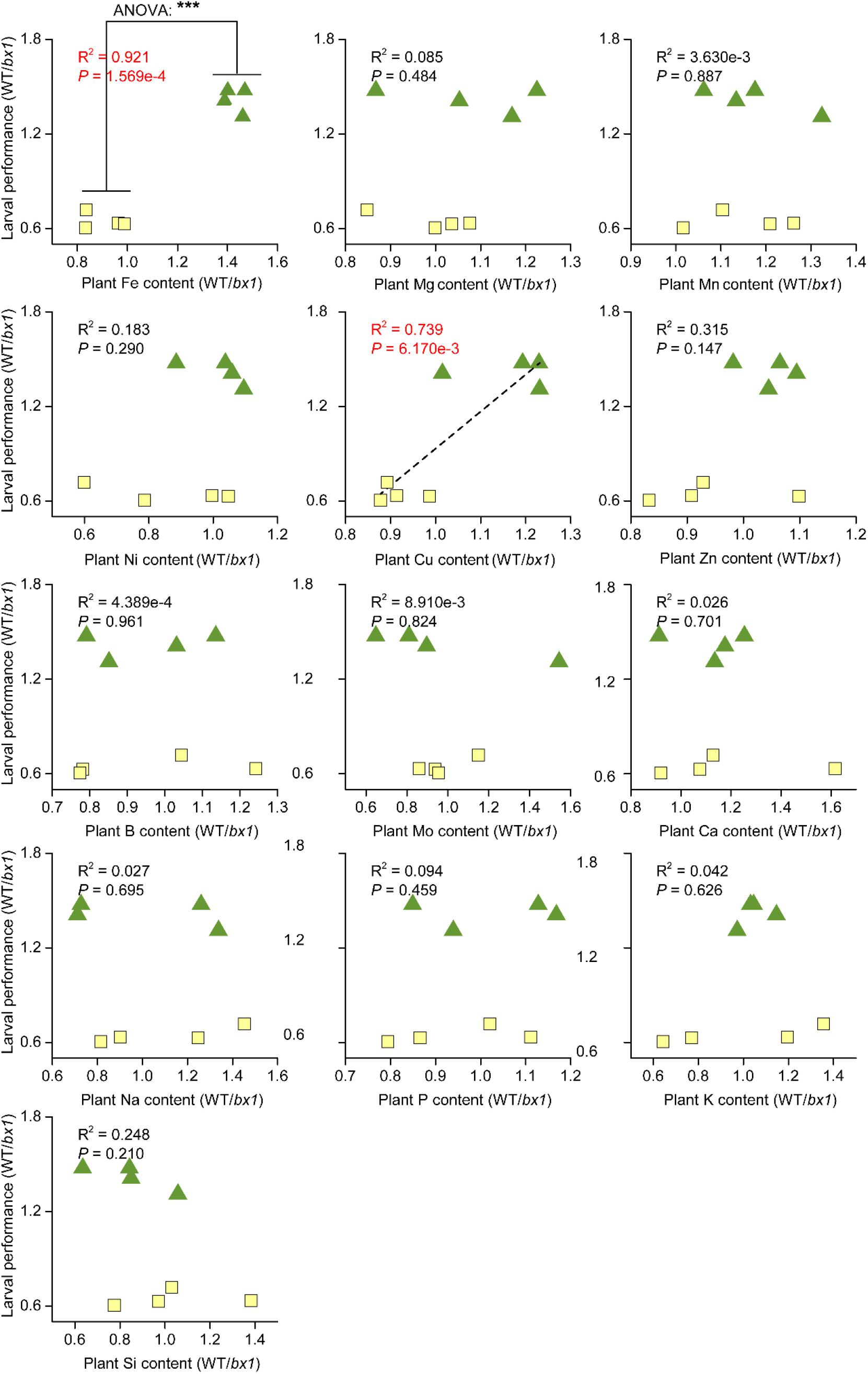
Correlations between benzoxazinoid-dependent larval performance and benzoxazinoid-dependent plant nutrients. Relative larval performance (WT/*bx1*) is correlated with relative leaf elemental concentrations (WT/*bx1*). Each point corresponds to data from one soil type. Green triangles correspond to soils where fall armyworm larvae grow better on WT than *bx1* mutant plants. Yellow squares correspond to soils where fall armyworm larvae grow better on *bx1* mutant than WT plants. R^2^ and *P* values of correlations are shown. Linear regression lines are drawn for significant linear correlations. Where clear grouping was observed (Fe), regression lines are omitted, and the *P*-value of an analysis of variance is shown instead \*\*\**p*<0.001).

**Fig. S6.**
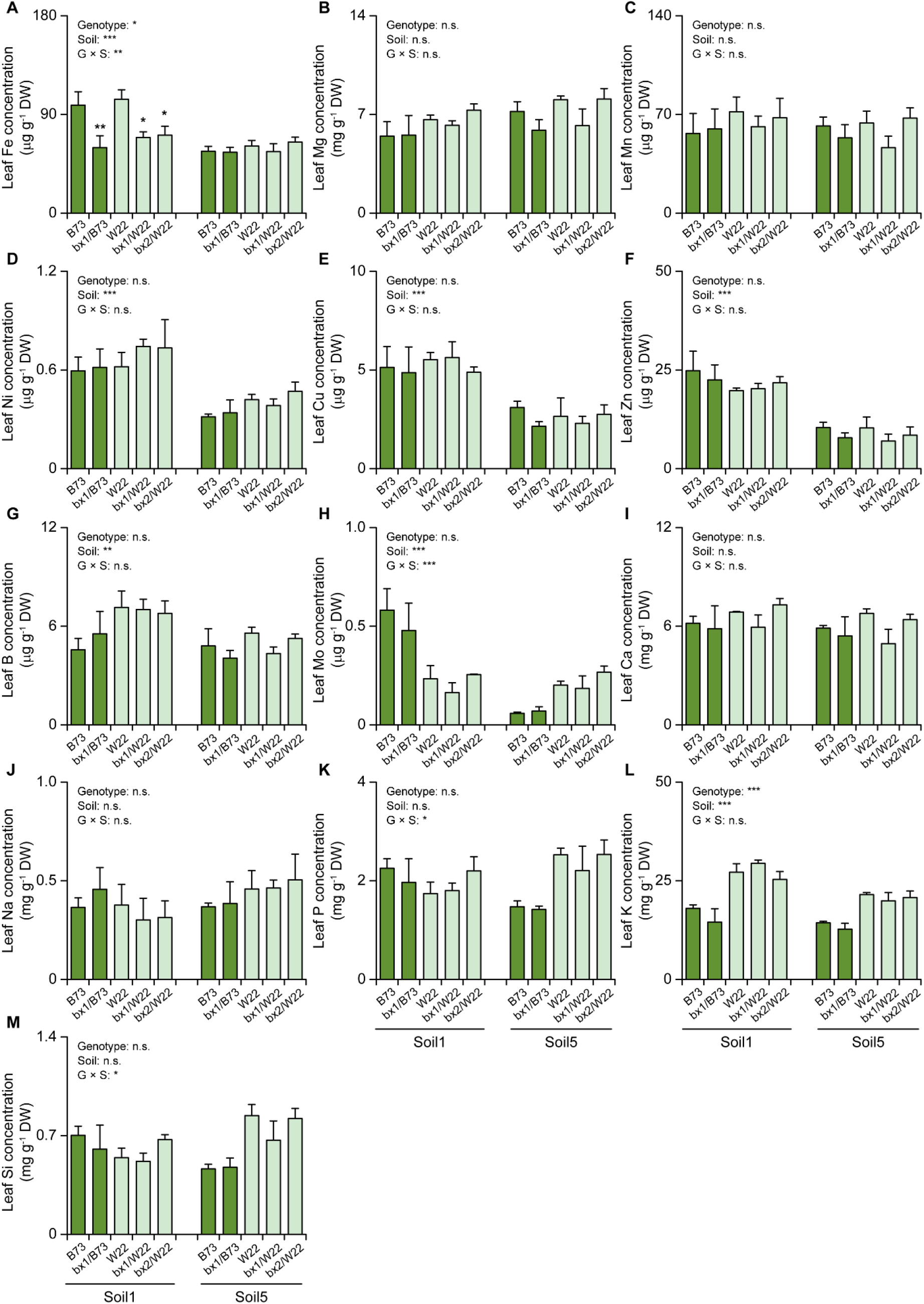
Leaf elemental analysis of *bx1* and *bx2* mutants in the W22 background. Concentrations of iron (Fe, A), magnesium (Mg, B), manganese (Mn, C), nickel (Ni, D), copper (Cu, E), zinc (Zn, F), boron (B, G), molydenum (Mo, H), calcium (Ca, I), sodium (Na, J), phosphorous (P, K), potassium (K, L), and silicon (Si, M) in leaves of *bx1*and *bx2* mutants as well as their corresponding wild type W22 grown in two soils (+SE, *n* = 3, with 3 to 4 individual plants pooled per replicate). DW, dry weight. Results of two-way ANOVAs are shown (n.s. not significant; \**P* < 0.05; \*\**P* < 0.01; \*\*\**P* < 0.001). Asterisks indicate significant differences between genotypes within the same soil (two-way ANOVA followed by pairwise comparisons through FDR-corrected LSMeans; \**P* < 0.05; \*\**P* < 0.01; \*\*\**P* < 0.001).

**Fig. S7.**
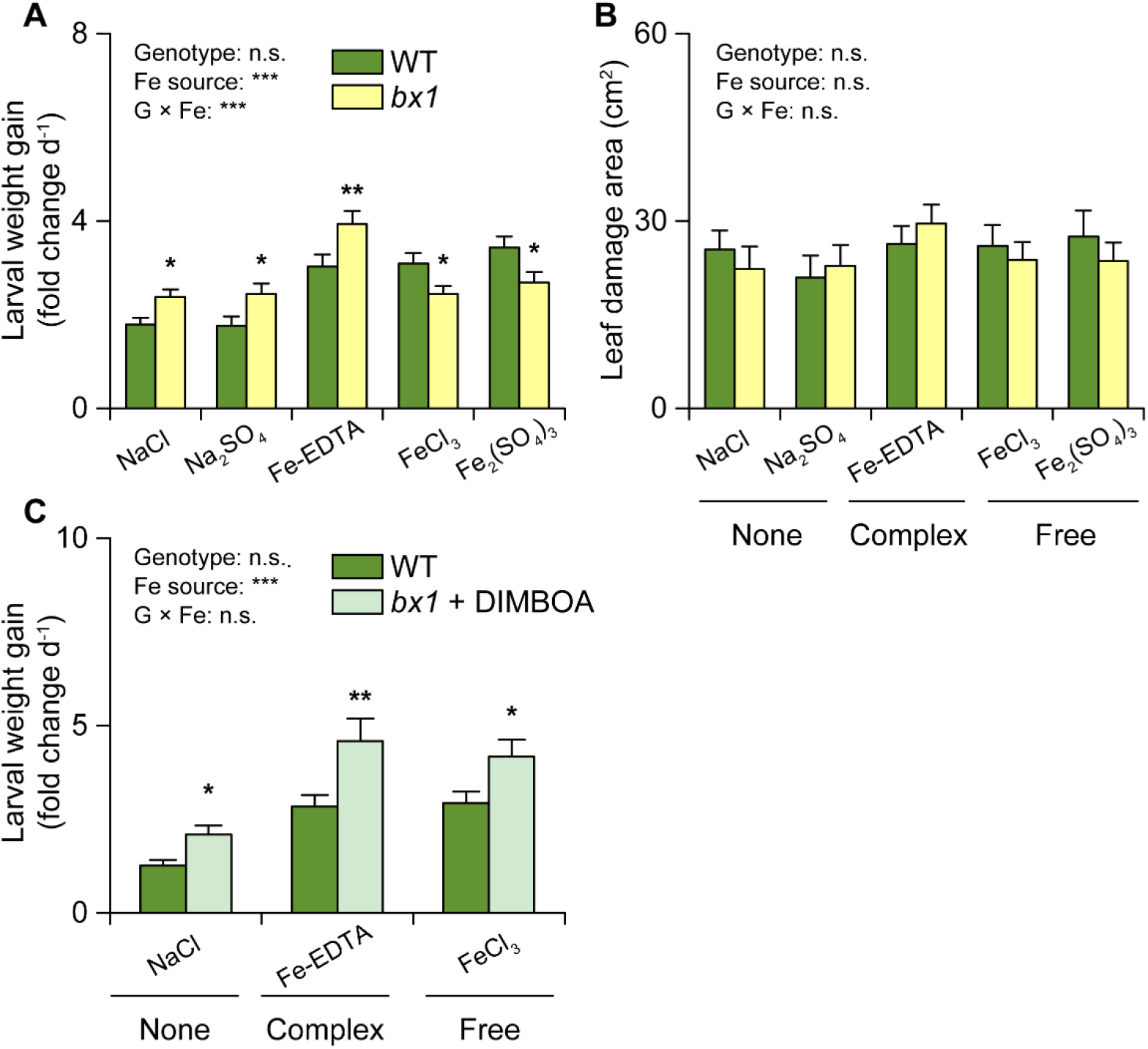
Soil-dependent benzoxazinoid resistance is driven by root iron supply. (A) Growth of *Spodoptera frugiperda* larvae feeding on wild type (WT) and *bx1* plants supplied with different iron sources (+SE, *n* = 14-15). (B) Consumed leaf area (+SE, *n* = 14-15). (C) Growth of *S. frugiperda* feeding on WT and *bx1* plants complemented with pure DIMBOA under different iron sources (+SE, *n* = 14-15). Results of two-way ANOVAs are shown (n.s. not significant; \*\*\**P* < 0.001). Asterisks indicate significant differences between genotypes within the same iron source (two-way ANOVA followed by pairwise comparisons through FDR-corrected LSMeans; \**P* < 0.05; \*\**P* < 0.01).

**Fig. S8.**
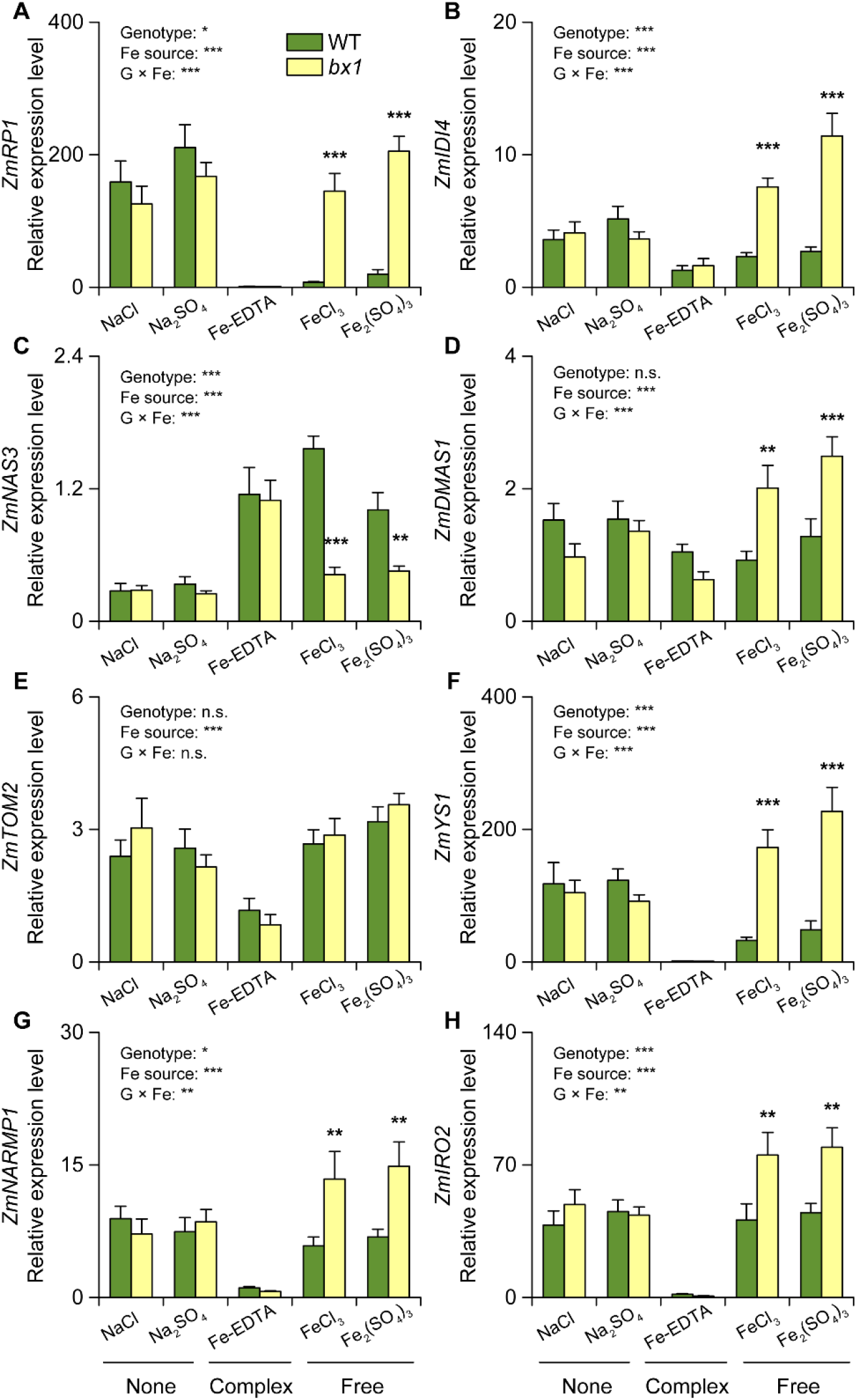
Interactions between root iron supply and benzoxazinoids determine leaf iron homeostasis. The expression levels of *ZmRP1*(A), *ZmIDI4* (B), *ZmNAS3* (C), *ZmDMAS1* (D), *ZmTOM2* (E), *ZmYS1* (F), *ZmNRAMP1* (G) and *ZmIRO2* (H) in wild type (WT) and *bx1* plants supplied with different iron sources (+SE, *n* = 14-15). Results of two-way ANOVAs are shown (n.s. not significant; \**P* < 0.05; \*\**P* < 0.01; \*\*\**P* < 0.001). Asterisks indicate significant differences within the same iron source (two-way ANOVA followed by pairwise comparisons through FDR-corrected LSMeans; \*\**P* < 0.01; \*\*\**P* < 0.001).

**Fig. S9.**
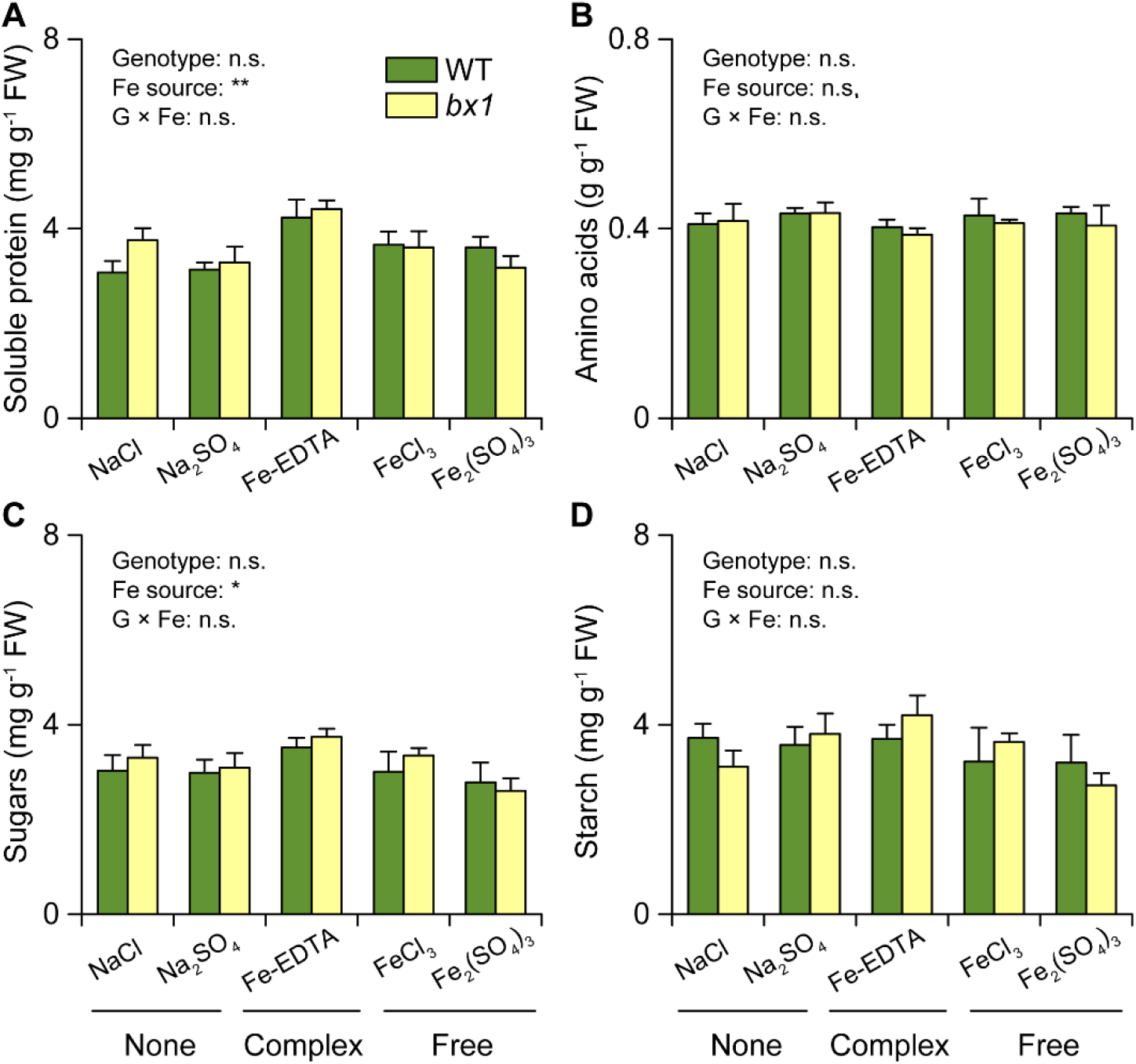
Changes in leaf herbivore performance are not explained by changes in leaf primary metabolism. Content of soluble protein (A), hydrolyzed amino acids (B), sugars (B) and starch (D) in wild type (WT) and *bx1* plants supplied with different iron sources (+SE, *n* = 14-15). Results of two-way ANOVAs are shown (n.s. not significant; \**P* < 0.05; \*\**P* < 0.01). No significant differences between genotypes within the same soil were observed (two-way ANOVA followed by pairwise comparisons through FDR-corrected LSMeans).

**Fig. S10.**
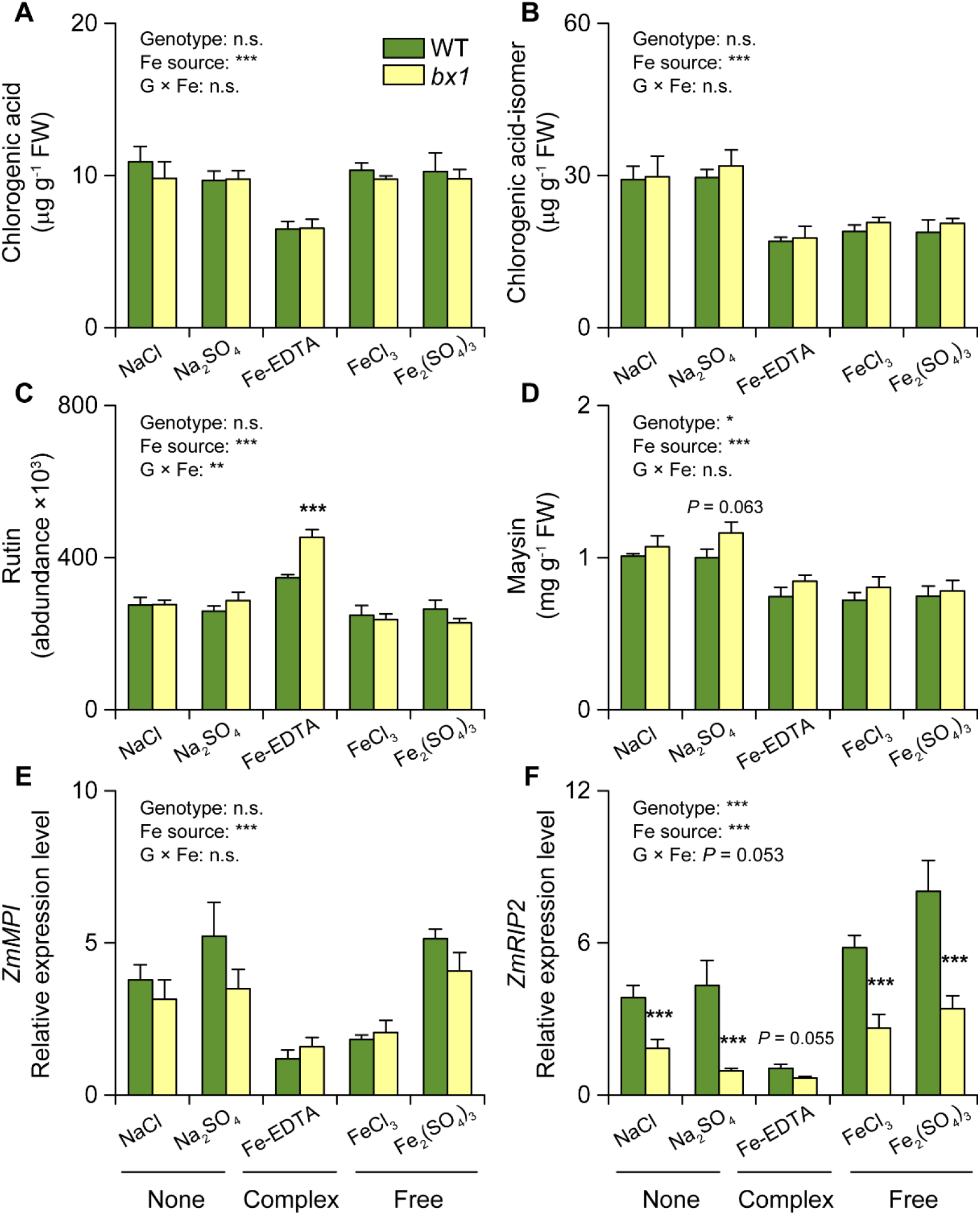
Soil iron and benzoxazinoids interactively reprogram a subset of leaf defenses. (A to D) Content of chlorogenic acid (A), chlorogenic acid-isomer (B), rutin (C) and maysin (D) in WT and *bx1* maize plants supplied with different iron sources (+SE, *n* = 8). FW, fresh weight. (E and F) Expression levels of *ZmMPI* (E) and *ZmRIP2* (F) in WT and *bx1* maize plants supplied with different iron sources (+SE, *n* = 8). Two-way ANOVA results testing for genotype and iron source effects are shown (n.s. not significant; \**P* < 0.05; \*\**P* < 0.01; \*\*\**P* < 0.001). Asterisks indicate significant differences between genotypes within the same soil (pairwise comparisons through FDR-corrected LSMeans; \*\*\**P* < 0.001).

**Fig. S11.**
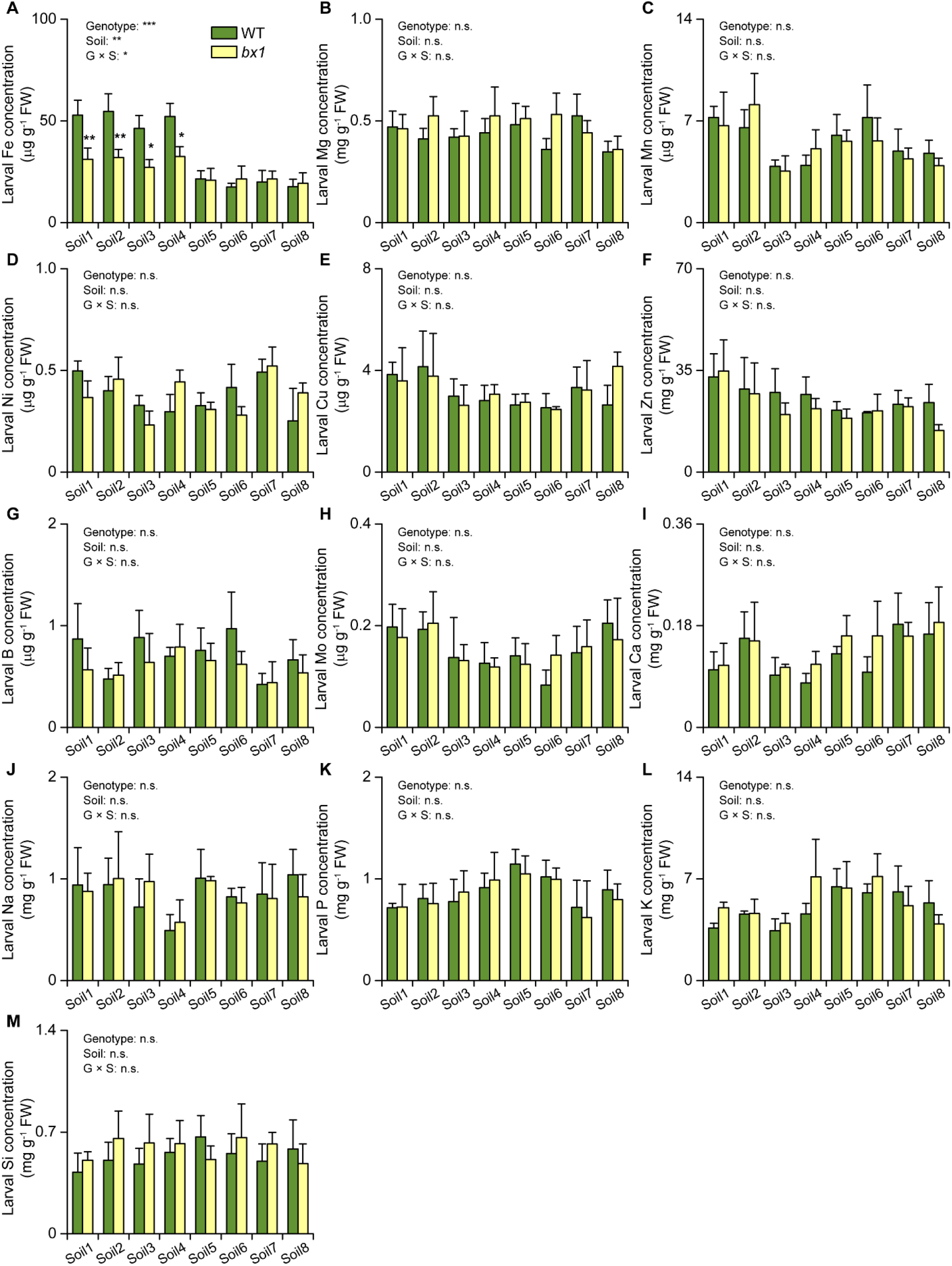
Elemental analysis of *Spodoptera frugiperda* larvae feeding on wild type (WT) and *bx1* plants. Concentrations of iron (Fe, A), magnesium (Mg, B), manganese (Mn, C), nickel (Ni, D), copper (Cu, E), zinc (Zn, F), boron (B, G), molydenum (Mo, H), calcium (Ca, I), sodium (Na, J), phosphorous (P, K), potassium (K, L), and silicon (Si, M) in *S. frugiperda* larvae feeding on wild type (WT) and *bx1* plants growing in different soils (+SE, *n* = 3, with 3 to 4 individual larvae pooled per replicate). FW, fresh weight. Results of two-way ANOVAs are shown (n.s. not significant; \**P* < 0.05; \*\**P* < 0.01; \*\*\**P* < 0.001). Asterisks indicate significant differences between genotypes within the same soil (two-way ANOVA followed by pairwise comparisons through FDR-corrected LSMeans; \**P* < 0.05; \*\**P* < 0.01).

**Fig. S12.**
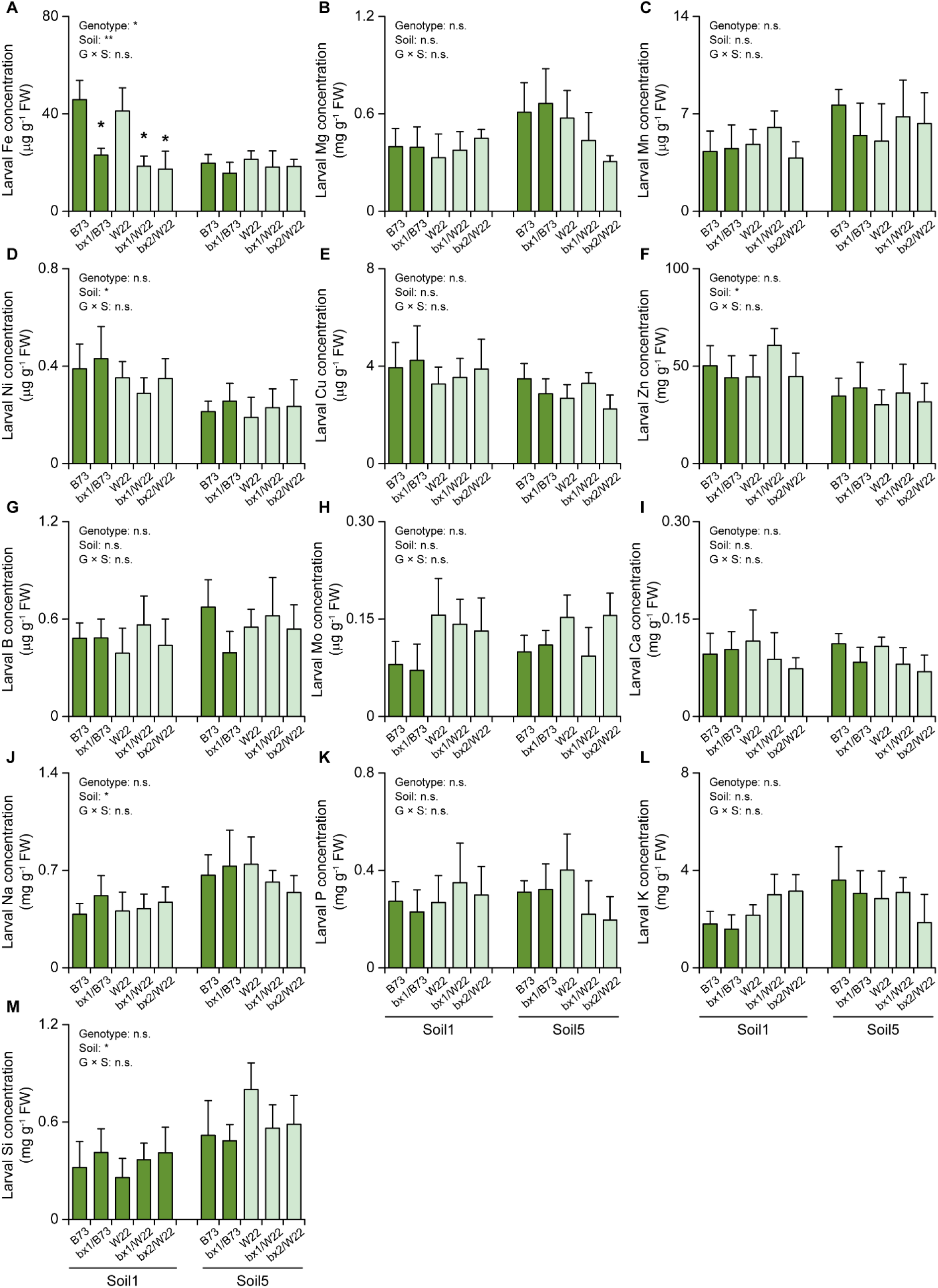
Elemental analysis of *Spodoptera frugiperda* larvae feeding on *bx1* and *bx2* mutants in the W22 background. Concentrations of iron (Fe, A), magnesium (Mg, B), manganese (Mn, C), nickel (Ni, D), copper (Cu, E), zinc (Zn, F), boron (B, G), molydenum (Mo, H), calcium (Ca, I), sodium (Na, J), phosphorous (P, K), potassium (K, L), and silicon (Si, M) in *S. frugiperda* larvae feeding *bx1* and *bx2* mutant plants in the W22 background grown in two different soils (+SE, *n* = 3, with 3 to 4 individual larvae pooled per replicate). FW, fresh weight. Results of two-way ANOVAs are shown (n.s. not significant; \**P* < 0.05; \*\**P* < 0.01). Asterisks indicate significant differences between genotypes within the same soil (two-way ANOVA followed by pairwise comparisons through FDR-corrected LSMeans; \**P* < 0.05).

**Fig. S13.**
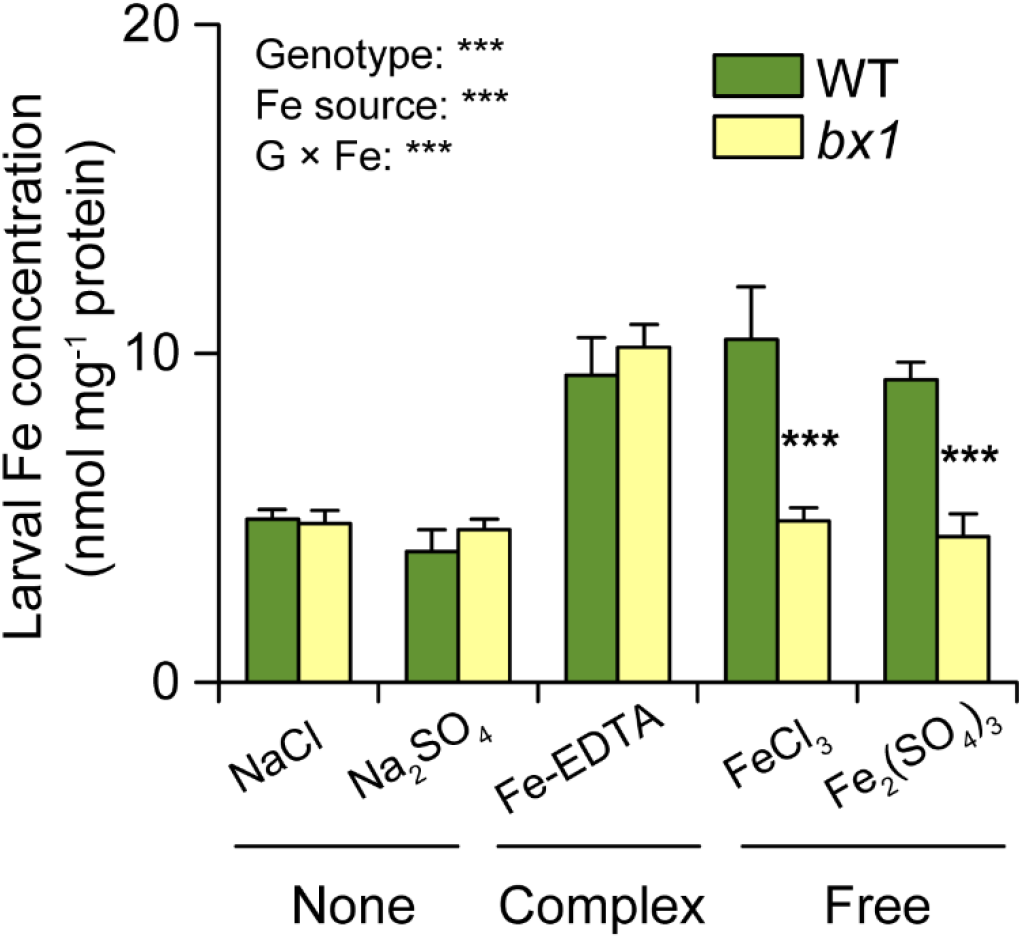
Influence of the interactions between benzoxazinoids and iron availability on larval iron levels. Iron contents of *Spodoptera frugiperda* larvae feeding on wild type (WT) and *bx1* plants (+SE, *n* = 5, with 3 individual larvae pooled per replicate). Results of two-way ANOVAs are shown (\*\*\**P* < 0.001). Asterisks indicate significant differences between genotypes within the same iron source (two-way ANOVA followed by pairwise comparisons through FDR-corrected LSMeans; \*\*\**P* < 0.001).

**Table S1.** Soil characteristics and classifications.

|  | **Latitude** | **Longitude** | **Soil types** | **Sand (%)** | **Silt (%)** | **Clay (%)** | **Cultivation genotypes** |
| --- | --- | --- | --- | --- | --- | --- | --- |
| **Soil1** | 31.42972 | 119.9155 | Anthrosol | 58 | 6 | 36 | B73, *bx1*/B73, W22, *bx1*/W22, and *bx2*/W22 |
| **Soil2** | 31.36122 | 119.6227 | Anthrosol | 50 | 8 | 42 | B73 and *bx1*/B73 |
| **Soil3** | 31.4873 | 119.9523 | Anthrosol | 36 | 12 | 52 | B73 and *bx1*/B73 |
| **Soil4** | 31.4666 | 119.8096 | Anthrosol | 48 | 6 | 46 | B73 and *bx1*/B73 |
| **Soil5** | 31.2994 | 119.5929 | Ferrosol | 38 | 6 | 56 | B73, *bx1*/B73, W22, *bx1*/W22, and *bx2*/W22 |
| **Soil6** | 31.3517 | 119.7333 | Ferrosol | 50 | 6 | 44 | B73 and *bx1*/B73 |
| **Soil7** | 31.17822 | 119.5458 | Ferrosol | 54 | 6 | 40 | B73 and *bx1*/B73 |
| **Soil8** | 31.2231 | 119.8401 | Ferrosol | 74 | 2 | 24 | B73 and *bx1*/B73 |

**Table S2.**
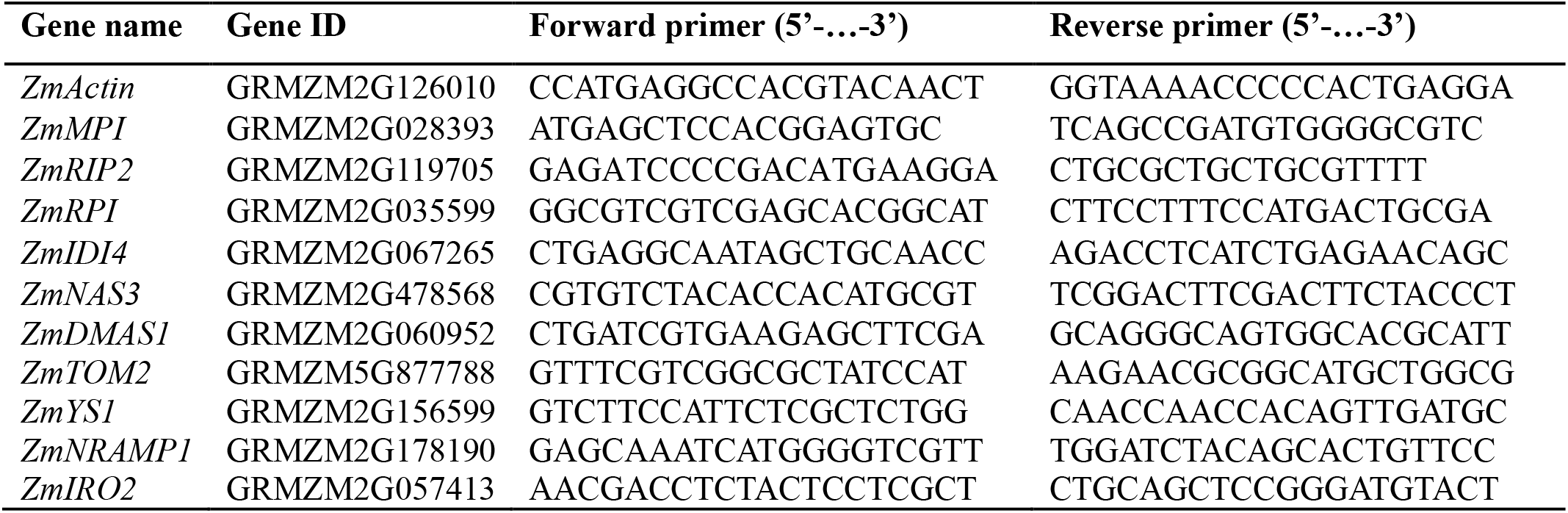
Primers used for QRT-PCR of target genes.

## References

1. S. C. Farina, E. A. Kane, L. P. Hernandez, Multifunctional structures and multistructural functions: Integration in the evolution of biomechanical systems. Integr. Comp. Biol. 59, 338–345 (2019).

2. D. S. Tawfik, Messy biology and the origins of evolutionary innovations. Nat. Chem. Biol. 6, 692–696 (2010).

3. R. J. Greenspan, The flexible genome. Nat. Rev. Genet. 2, 383–387 (2001).

4. E. H. Neilson, J. Q. D. Goodger, I. E. Woodrow, B. L. Møller, Plant chemical defense: at what cost? Trends Plant Sci. 18, 250–258 (2013).

5. N. R. Friedman et al., Evolution of a multifunctional trait: shared effects of foraging ecology and thermoregulation on beak morphology, with consequences for song evolution. Proc. Roy. Soc. B-Biol. Sci. 286 (2019).

6. B. Li et al., Convergent evolution of a metabolic switch between aphid and caterpillar resistance in cereals. Sci. Adv. 4, eaat6797 (2018).

7. R. Li et al., Prioritizing plant defence over growth through WRKY regulation facilitates infestation by non-target herbivores. eLife 4, e04805 (2015).

8. T. Hartmann, From waste products to ecochemicals: fifty years research of plant secondary metabolism. Phytochemistry 68, 2831–2846 (2007).

9. A. Mithofer, W. Boland, Plant defense against herbivores: Chemical aspects. Annu. Rev. Plant Biol. 63, 431–450 (2012).

10. N. K. Clay, A. M. Adio, C. Denoux, G. Jander, F. M. Ausubel, Glucosinolate metabolites required for an Arabidopsis innate immune response. Science 323, 95–101 (2009).

11. G. Agati, M. Tattini, Multiple functional roles of flavonoids in photoprotection. New Phytol. 186, 786–793 (2010).

12. M. Huang et al., The major volatile organic compound emitted from *Arabidopsis thaliana* flowers, the sesquiterpene (*E*)-beta-caryophyllene, is a defense against a bacterial pathogen. New Phytol. 193, 997–1008 (2012).

13. N. B. Schmid et al., Feruloyl-coa 6′-hydroxylase1-dependent coumarins mediate iron acquisition from alkaline substrates in Arabidopsis. Plant Physiol. 164, 160–172 (2014).

14. E. Soubeyrand et al., The peroxidative cleavage of kaempferol contributes to the biosynthesis of the benzenoid moiety of ubiquinone in plants. Plant Cell 30, 2910–2921 (2018).

15. M. Erb, D. J. Kliebenstein, Plant secondary metabolites as defenses, regulators, and primary metabolites: The blurred functional trichotomy. Plant Physiol. 184, 39–52 (2020).

16. E. Pichersky, R. A. Raguso, Why do plants produce so many terpenoid compounds? New Phytol. 220, 692–702 (2018).

17. D. J. Ballhorn, A. Pietrowski, R. Lieberei, Direct trade-off between cyanogenesis and resistance to a fungal pathogen in lima bean (*Phaseolus lunatus* L.). J. Ecol. 98, 226–236 (2010).

18. D. Baek, H. J. Chun, D. J. Yun, M. C. Kim, Cross-talk between phosphate starvation and other environmental stress signaling pathways in plants. Mol. Cells 40, 697–705 (2017).

19. L. A. J. Mur, C. Simpson, A. Kumari, A. K. Gupta, K. J. Gupta, Moving nitrogen to the centre of plant defence against pathogens. Ann. Bot. 119, 703–709 (2017).

20. A. Martinez-Medina, S. C. M. Van Wees, C. M. J. Pieterse, Airborne signals from Trichoderma fungi stimulate iron uptake responses in roots resulting in priming of jasmonic acid-dependent defences in shoots of *Arabidopsis thaliana* and *Solanum lycopersicum*. Plant Cell Environ. 40, 2691–2705 (2017).

21. I. A. Stringlis et al., MYB72-dependent coumarin exudation shapes root microbiome assembly to promote plant health. Proc. Natl. Acad. Sci. U.S.A. 115, 5213–5222 (2018).

22. J. D. Schade, M. Kyle, S. E. Hobbie, W. F. Fagan, J. J. Elser, Stoichiometric tracking of soil nutrients by a desert insect herbivore. Ecol. Lett. 6, 96–101 (2003).

23. M. Frey, K. Schullehner, R. Dick, A. Fiesselmann, A. Gierl, Benzoxazinoid biosynthesis, a model for evolution of secondary metabolic pathways in plants. Phytochemistry 70, 1645–1651 (2009).

24. H. M. Niemeyer, Hydroxamic acids derived from 2-hydroxy-2H-1,4-benzoxazin-3(4H)-one: Key defense chemicals of cereals. J. Agric. Food Chem. 57, 1677–1696 (2009).

25. S. Ahmad et al., Benzoxazinoid metabolites regulate innate immunity against aphids and fungi in maize. Plant Physiol. 157, 317–327 (2011).

26. L. N. Meihls et al., Natural variation in maize aphid resistance is associated with 2,4-dihydroxy-7-methoxy-1,4-benzoxazin-3-one glucoside methyltransferase activity. Plant Cell 25, 2341–2355 (2013).

27. C. A. M. Robert et al., A specialist root herbivore exploits defensive metabolites to locate nutritious tissues. Ecol. Lett. 15, 55–64 (2012).

28. L. Bigler, A. Baumeler, C. Werner, M. Hesse, Detection of noncovalent complexes of hydroxamic-acid derivatives by means of electrospray mass spectrometry. Helv. Chim. Acta 79, 1701–1709 (1996).

29. L. Hu et al., Plant iron acquisition strategy exploited by an insect herbivore. Science 361, 694–697 (2018).

30. G. Glauser et al., Induction and detoxification of maize 1,4-benzoxazin-3-ones by insect herbivores. Plant J. 68, 901–911 (2011).

31. D. Maag et al., Highly localized and persistent induction of *Bx1*-dependent herbivore resistance factors in maize. Plant J. 88, 976–991 (2016).

32. C. Poschenrieder, R. P. Tolra, J. Barcelo, A role for cyclic hydroxamates in aluminium resistance in maize? J. Inorg. Biochem. 99, 1830–1836 (2005).

33. C. Curie et al., Maize *yellow stripe1* encodes a membrane protein directly involved in Fe(III) uptake. Nature 409, 346–349 (2001).

34. S. N. Chorianopoulou, Y. I. Saridis, M. Dimou, P. Katinakis, D. L. Bouranis, Arbuscular mycorrhizal symbiosis alters the expression patterns of three key iron homeostasis genes, ZmNAS1, ZmNAS3, and ZmYS1, in S deprived maize plants. Front. Plant Sci. 6, 257 (2015).

35. T. Nozoye, H. Nakanishi, N. K. Nishizawa, Characterizing the crucial components of iron homeostasis in the maize mutants *ys1* and *ys3*. PLoS One 8, e62567 (2013).

36. M. Jahangir, I. B. Abdel-Farid, Y. H. Choi, R. Verpoorte, Metal ion-inducing metabolite accumulation in *Brassica rapa*. J. Plant Physiol. 165, 1429–1437 (2008).

37. G. Vigani et al., Knocking down mitochondrial iron transporter (MIT) reprograms primary and secondary metabolism in rice plants. J. Exp. Bot. 67, 1357–1368 (2016).

38. J. H. Herlihy, T. A. Long, J. M. McDowell, Iron homeostasis and plant immune responses: Recent insights and translational implications. J. Biol. Chem. 295, 13444–13457 (2020).

39. B. R. Wiseman, M. E. Snook, R. L. Wilson, D. J. Isenhour, Allelochemical content of selected popcorn silks: Effects on growth of corn earworm larvae (lepidoptera: Noctuidae). J. Econ. Entomol. 85, 2500–2504 (1992).

40. M. C. Tamayo, M. Rufat, J. M. Bravo, B. San Segundo, Accumulation of a maize proteinase inhibitor in response to wounding and insect feeding, and characterization of its activity toward digestive proteinases of *Spodoptera littoralis* larvae. Planta 211, 62–71 (2000).

41. W. P. Chuang et al., Caterpillar attack triggers accumulation of the toxic maize protein RIP2. New Phytol. 201, 928–939 (2014).

42. N. Vonwiren, S. Mori, H. Marschner, V. Romheld, Iron inefficiency in maize mutant ys1 (*Zea mays* L cv yellow-stripe) is caused by a defect in uptake of iron phytosiderophores. Plant Physiol. 106, 71–77 (1994).

43. L. Sack, T. N. Buckley, Trait multi-functionality in plant stress response. Integr. Comp. Biol. 60, 98–112 (2020).

44. S. Zhou, A. Richter, G. Jander, Beyond defense: Multiple functions of benzoxazinoids in maize metabolism. Plant Cell Physiol. 10.1093/pcp/pcy064, 1528–1537 (2018).

45. D. J. Kliebenstein, Plant nutrient acquisition entices herbivore. Science 361, 642–643 (2018).

46. F. C. Wouters, B. Blanchette, J. Gershenzon, D. G. Vassao, Plant defense and herbivore counter-defense: benzoxazinoids and insect herbivores. Phytochem. Rev. 15, 1127–1151 (2016).

47. F. C. Wouters et al., Reglucosylation of the benzoxazinoid DIMBOA with inversion of stereochemical configuration is a detoxification strategy in lepidopteran herbivores. Angew. Chem. Int. Edit. 53, 11320–11324 (2014).

48. A. Kohler et al., Within-plant distribution of 1,4-benzoxazin-3-ones contributes to herbivore niche differentiation in maize. Plant Cell Environ. 38, 1081–1093 (2014).

49. L. Hu et al., Root exudate metabolites drive plant-soil feedbacks on growth and defense by shaping the rhizosphere microbiota. Nat. Commun. 9, 2738 (2018).

50. Y. Lou, I. T. Baldwin, Nitrogen supply influences herbivore-induced direct and indirect defenses and transcriptional responses in *Nicotiana attenuata*. Plant Physiol. 135, 496–506 (2004).

51. T. J. Massad, L. A. Dyer, C. G. Vega, Costs of defense and a test of the carbon-nutrient balance and growth-differentiation balance hypotheses for two co-occurring classes of plant defense. PLoS One 7, e47554 (2012).

52. W. C. Wetzel, H. M. Kharouba, M. Robinson, M. Holyoak, R. Karban, Variability in plant nutrients reduces insect herbivore performance. Nature 539, 425–427 (2016).

53. D. Debona, F. A. Rodrigues, L. E. Datnoff, Silicon’s role in abiotic and biotic plant stresses. Annu. Rev. Phytopathol. 55, 85–107 (2017).

54. C. Cabot et al., A role for zinc in plant defense against pathogens and herbivores. Front. Plant Sci. 10, 1171 (2019).

55. E. H. Verbon et al., Iron and Immunity. Annu. Rev. Phytopathol. 55, 355–375 (2017).

56. E. N. Kudjordjie, R. Sapkota, S. K. Steffensen, I. S. Fomsgaard, M. Nicolaisen, Maize synthesized benzoxazinoids affect the host associated microbiome. Microbiome 7 (2019).

57. T. E. A. Cotton et al., Metabolic regulation of the maize rhizobiome by benzoxazinoids. ISME J. 13, 1647–1658 (2019).

58. E. S. Bakker, M. E. Ritchie, H. Olff, D. G. Milchunas, J. M. Knops, Herbivore impact on grassland plant diversity depends on habitat productivity and herbivore size. Ecol. Lett. 9, 780–788 (2006).

59. T. Ohgushi, Eco-evolutionary dynamics of plant-herbivore communities: incorporating plant phenotypic plasticity. Curr. Opin. Insect Sci. 14, 40–45 (2016).

60. D. A. Wardle et al., Ecological linkages between aboveground and belowground biota. Science 304, 1629–1633 (2004).

61. V. Tzin et al., Dynamic maize responses to aphid feeding are revealed by a time series of transcriptomic and metabolomic assays. Plant Physiol. 169, 1727–1743 (2015).

62. D. Maag et al., 3-*β*-D-Glucopyranosyl-6-methoxy-2-benzoxazolinone (MBOA-*N*-Glc) is an insect detoxification product of maize 1,4-benzoxazin-3-ones. Phytochemistry 102, 97–105 (2014).

63. H. Yuan et al., Warming facilitates microbial reduction and release of arsenic in flooded paddy soil and arsenic accumulation in rice grains. J. Hazard. Mater. 408, 124913 (2021).

64. N. M. van Dam, M. Horn, M. Mares, I. T. Baldwin, Ontogeny constrains systemic protease inhibitor response in *Nicotiana attenuata*. J. Chem. Ecol. 27, 547–568 (2001).

65. T. Docimo et al., The first step in the biosynthesis of cocaine in *Erythroxylum coca*: the characterization of arginine and ornithine decarboxylases. Plant Mol. Biol. 78, 599–615 (2012).

66. R. A. R. Machado et al., Leaf-herbivore attack reduces carbon reserves and regrowth from the roots via jasmonate and auxin signaling. New Phytol. 200, 1234–1246 (2013).

67. D. Maag et al., Maize domestication and anti-herbivore defences: Leaf-specific dynamics during early ontogeny of maize and its wild ancestors. PLoS One 10, e0135722 (2015).

68. M. L. Wong, J. F. Medrano, Real-time PCR for mRNA quantitation. Biotechniques 39, 75–85 (2005).

69. F. Missirlis et al., Characterization of mitochondrial ferritin in *Drosophila*. Proc. Natl. Acad. Sci. U.S.A. 103, 5893–5898 (2006).

70. R. A. R. Machado, C. C. M. Arce, A. P. Ferrieri, I. T. Baldwin, M. Erb, Jasmonate-dependent depletion of soluble sugars compromises plant resistance to *Manduca sexta*. New Phytol. 207, 91–105 (2015).

71. Y. Benjamini, Y. Hochberg, Controlling the false discovery rate: a practical and powerful approach to multiple testing. J. R. Stat. Soc. Ser. B-Stat. Methodol. 57, 289–300 (1995).

72. M. R. Herve, RVAideMemoire: Testing and plotting procedures for biostatistics. Available at. https://cran.r-project.org/web/packages/RVAideMemoire/index.html. (2021).

73. J. Oksanen et al., Vegan: community ecology package. R package version 2.0-10 (The Comprehensive R Archive Network (CRAN), Vienna, Austria. 2013). Available at. http://CRAN.R-project.org/package=vegan. (2013).

74. D. Bates, M. Machler, B. M. Bolker, S. C. Walker, Fitting linear mixed-effects models using lme4. J. Stat. Softw. 67, 1–48 (2015).

75. R. V. Lenth, Least-squares means: The R Package lsmeans. J. Stat. Softw. 69, 1–33 (2016).

